# Senotherapeutic peptide reduces skin biological age and improves skin health markers

**DOI:** 10.1101/2020.10.30.362822

**Authors:** Alessandra Zonari, Lear E. Brace, Kallie Z. Al-Katib, William F. Porto, Daniel Foyt, Mylieneth Guiang, Edgar Andres Ochoa Cruz, Bailey Marshall, Willian G. Salgueiro, Mehmet Dinçer Inan, Mizanur Rahman, Taslim Anupom, Siva Vanapalli, Marcelo A. Mori, Octavio L. Franco, Carolina R. Oliveira, Mariana Boroni, Juliana L. Carvalho

## Abstract

Skin aging has been primarily related to aesthetics and beauty. Therefore, interventions have focused on reestablishing skin appearance, but not necessarily skin health, function, and resilience. Recently, cellular senescence was shown to play a role in age-related skin function deterioration and influence organismal health and, potentially, longevity. In the present study, a two-step screening was performed to identify peptides capable of reducing cellular senescence in human dermal fibroblasts (HDF) from Hutchinson-Gilford Progeria (HGPS) patients. From the top four peptides of the first round of screening, we built a 764-peptide library using amino acid scanning, of which the second screen led to the identification of peptide 14. Peptide 14 effectively decreased HDF senescence induced by HGPS, chronological aging, ultraviolet-B radiation, and etoposide treatment, without inducing significant cell death, and likely by modulating longevity and senescence pathways. We further validated the effectiveness of peptide 14 using human skin equivalents and skin biopsies, where peptide 14 promoted skin health and reduced senescent cell markers, as well as the biological age of samples, according to the Skin-Specific DNA methylation clock, MolClock. Topical application of peptide 14 outperformed Retinol treatment, the current gold-standard in “anti-aging” skin care. Finally, we determined that peptide 14 is safe for long-term applications and also significantly extends both the lifespan and healthspan of *C. elegans* worms tested in two independent testings. This highlights the potential for geroprotective applications of the senotherapeutic compounds identified using our screening platform beyond the skin.

## Introduction

Aging is an intrinsic process associated with tissue dysfunction, loss of health, and increased rates of degenerative diseases^1^. In the skin, aging is visually perceived by the accumulation of wrinkles, loss of skin elasticity, and aberrant pigmentation^2, 3^ Many intrinsic and extrinsic processes underlying skin aging have already been described. For instance, intrinsic hormonal changes associated with aging lead to lower deposition of hyaluronic acid and collagen by fibroblasts and reduced cellular proliferation, which compromises cell turnover in skin^2,4^ Extrinsic factors such as sun exposure and smoking have also been shown to lead to the accumulation of inflammatory signals and decreased vascularization in the skin, resulting in altered cellular metabolism, and melanin synthesis^2,4^. The understanding of these and other events that influence skin aging have paved the way for sunscreens, antioxidants, collagen and hyaluronic acid inducers, lightening and anti-inflammatory agents for skin care^2^.

The presence and accumulation of senescent cells with age in the skin has been long observed^5–8^. Senescent cells were previously considered to only be byproducts of tissue aging, but current studies show them to be active inducers of aging and dysfunction^9–11^, justifying the need for the development of senotherapeutic approaches to treat aging-related diseases^12-14^. Considering that senescent cells play an active role in skin aging, a mosaic model has been proposed to explain age-related skin deterioration, in which senescent cells are induced by both intrinsic and extrinsic stimuli, such as chronological age, genetic background, and environmental conditions^15,16^. According to this model, the aforementioned stimuli synergize and lead to senescent cell accumulation in the skin, which actively promotes further tissue dysfunction, affecting the local tissue microenvironment through the senescence-associated secretory phenotype (SASP). The SASP includes several proinflammatory molecules^16,17^, and compromises epidermal stem cell renewal^18^, ECM deposition, melanin synthesis, among others^19^. In this scenario, the selective elimination or suppression of senescent cell populations might effectively interrupt such feedforward dynamics in the skin by terminating the SASP. This strategy has been tested using animal models in other tissue niches, such as the hematopoietic system^12^, bone^20^, hair, and the overall organism^13^. Currently, evidence supporting the hypothesis that senotherapeutic strategies may be effective at promoting human skin rejuvenation is limited to one study which documented the positive effects of topical Rapamycin treatment in the skin^21^. Therefore, the knowledge regarding cellular senescence has not been definitively translated into effective interventions for skin aging.

Here, we screened novel peptides according to their potential to significantly reduce cellular senescence levels in a HGPS model of cellular aging. One peptide obtained from the screening and optimization process, peptide 14, was chosen for further experiments. Peptide 14 decreased the level of cellular senescence in many experimental models, promoting tissue rejuvenation in both 2D and 3D systems, as measured by cellular and molecular markers of aging, senescence, and DNA methylation (DNAm). Furthermore, compared to Retinol, the gold-standard for “antiaging” skin treatments, human skins treated with peptide 14 presented increased epidermal thickness and showed improvement in numerous skin health markers. Importantly, peptide 14 was shown to be safe in long-term toxicology experiments and also to significantly increase the median and maximum lifespan of *C. elegans* worms using two different culture environments (agar plates and microfluidic devices), indicating its potential as a geroprotective agent. Taken together, we suggest that senotherapeutic molecules may contribute to the development of the next generation of skincare products that are specifically designed to intervene with underlying causes of skin aging and promote skin health on a molecular, cellular and tissue level.

## Results

### Screening of a peptide library identifies new senotherapeutic compounds

A library of 164 non-cytotoxic peptides from previous work^22^, was synthesized and then screened for the presence of senotherapeutic molecules by measuring senescence-associated betagalactosidase staining (SA-BGal) of primary human dermal fibroblasts (HDF) from HGPS patients, known to have a significant population of senescent cells (Sup. Fig. 1A). The top four performing peptides were further validated (Sup. Fig. 1B) and used as reference amino acid sequences to generate 764 leads. The new peptide library was then assessed for toxicity and senotherapeutic potential (Fig. 1A). Five peptides displaying the lowest cytotoxicity and highest senotherapeutic potential were validated in additional experimental replicates, and peptide 14 was selected for further investigation (Fig 1B, C). HGPS-HDFs treated with peptide 14 displayed a significant decrease in the mRNA expression of genes related to aging, inflammation and SASP (P16, P21 and IL-8) (Fig 1D).

**Figure 1.**
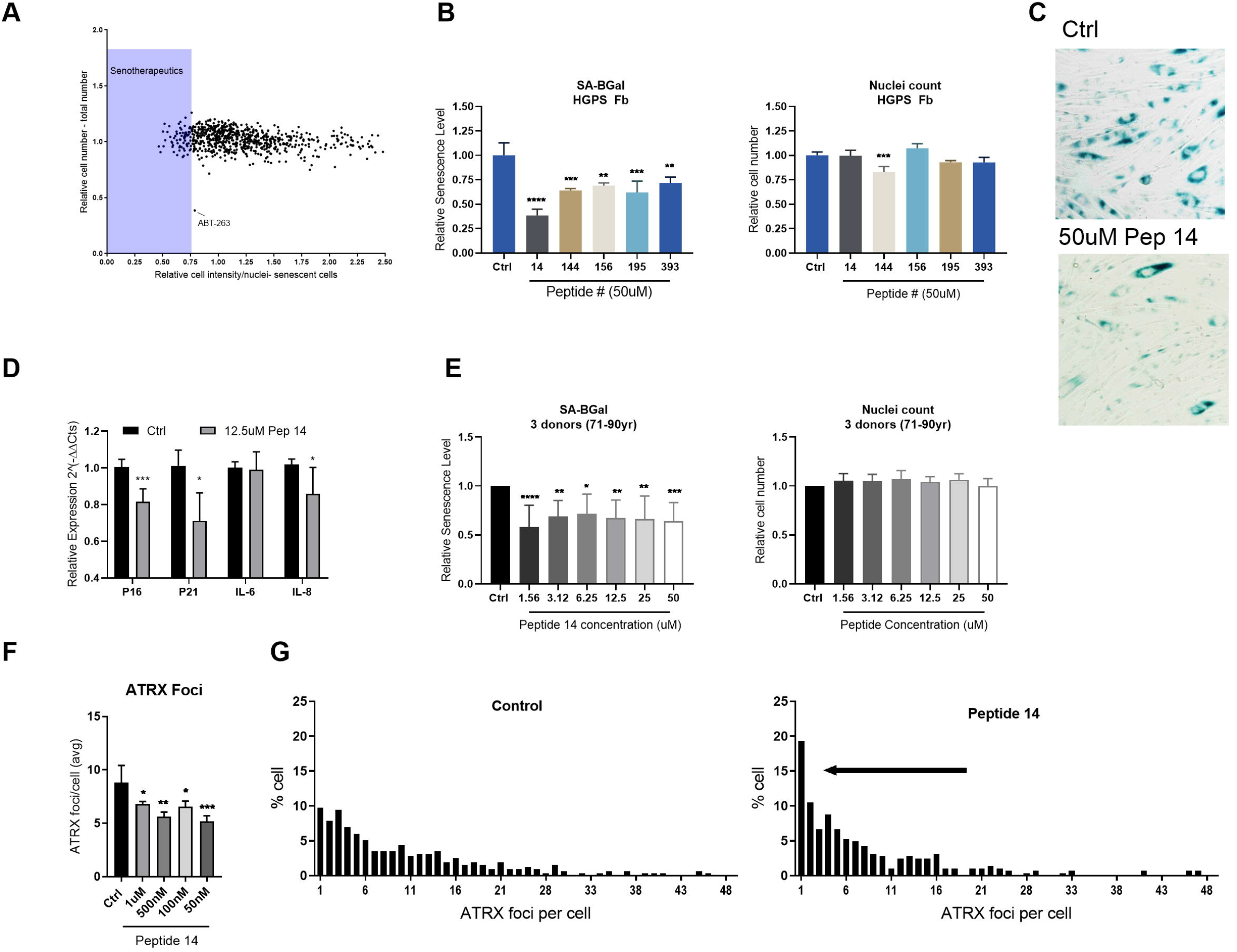
Screening of a peptide library identifies new senotherapeutic compounds. (A) Screening of the 764-peptide leads according to the senotherapeutic potential. Peptides that reduced the number of senescent HDFs isolated from HGPS patients by more than 25% were considered senotherapeutics and highlighted in the blue shaded area. (B) Relative cellular senescence and cell number of HGPS HDFs treated with 50 μM of the top 5 peptides screened in A. (C) Representative SA-BGal stained image of HGPS HDFs treated with 50 μM peptide 14 (Pep 14) compared to untreated controls. (D) mRNA expression of HGPS fibroblasts treated with 12.5 μM peptide 14 for 48 hr. (E) Relative cellular senescence and cell number of HDFs to non-treated controls obtained from 3 healthy donors with ages between 71 and 91 years and treated with different concentrations of peptide 14. (F) Mean ATRX foci/cell of 84 yr HDFs treated with different concentrations of peptide 14. (G) Representative analysis of the percentage of 84yr HDFs with specific numbers of ATRX foci/cell in control and 500 nM peptide 14-treated samples. The black arrow indicates that peptide 14-treated cells presented lower numbers of ATRX foci/cell, compared to untreated controls. Data are representative of ≥3 independent experiments and were analyzed using student’s t-test. *p<0.05; **p<0.01; ***p<0.001; ****p<0.0001, compared to untreated controls.

As it is well known that senescent cell populations increase with age, we assessed whether peptide 14 would also be effective in reducing the senescence levels of HDF obtained from healthy aged donors. Increased SA-BGal positive cells were observed with age of HDF, as was the accumulation of nuclear ATRX foci, an early marker of cellular senescence (Sup. Fig. 1C,D)^23^. We observed that the relative number of SA-BGal positive cells decreased with numerous concentrations of peptide 14 treatment without altering overall cell number (Fig. 1E). Aged HDF treated with 50 nM to 1 μM peptide 14 led to a statistically significant decrease in the average ATRX foci/nuclei (Fig. 1F), corroborating SA-BGal results. The distribution of ATRX foci per nuclei increased with age (Sup. Fig. 1E) and treatment of 84yr HDF with 500 nM peptide 14 also shifted the average number of foci per cell closer to a younger ATRX distribution profile (Fig. 1G). In sum, peptide 14 reduces cellular senescence in both highly senescent HGPS HDFs and age-accumulated senescent cells in primary HDFs from different donors.

### Peptide 14 protects fibroblasts from acute UVB and etoposide-induced cellular senescence

Ultraviolet-B (UVB) exposure is recognized as a relevant driver of skin aging^4,24–26^ and is also an effective strategy to induce premature cellular senescence *in vitro*^27–30^. To determine whether peptide 14 could also protect skin cells from extrinsic inducers of cellular senescence, we treated UVB-exposed HDF with peptide 14 and assessed cellular senescence markers. UVB doses of 0.05 J/cm^2^ and 0.1 J/cm^2^ effectively induced dose-dependent cellular senescence, as verified by SA-BGal staining relative to cell count, and reduced overall cell count as expected (Fig. 2A). UVB-exposed fibroblasts from 30 and 79 year old donors treated with increasing concentrations of peptide 14 presented decreased SA-BGal staining/nuclei and prevented UVB-induced cell death (Fig. 2B,C). To support the reduction in senescence we measured gene expression of cells exposed to 0.1 J/cm^2^ and treated with 500 nM peptide 14. UVB-damaged 30yr fibroblasts treated with peptide 14 displayed a significant decrease in the mRNA expression of genes related to aging, inflammation and SASP (B2M, IL-6, and IL-8), and a trend towards a decrease in senescence (P16) (Fig. 2D). In UVB-dosed 79yr HDFs treated with peptide 14, a trend was observed for decreased gene expression of P16 and B2M but no change in inflammatory markers was observed with peptide 14 treatment (Fig. 2D). On the protein level, peptide 14 also decreased the levels of the senescence marker P21 in 30yr and 79yr HDFs after treatment with an acute dose of UVB (Fig. 2E, representative image, F, 3-4 independent experiments).

**Figure 2.**
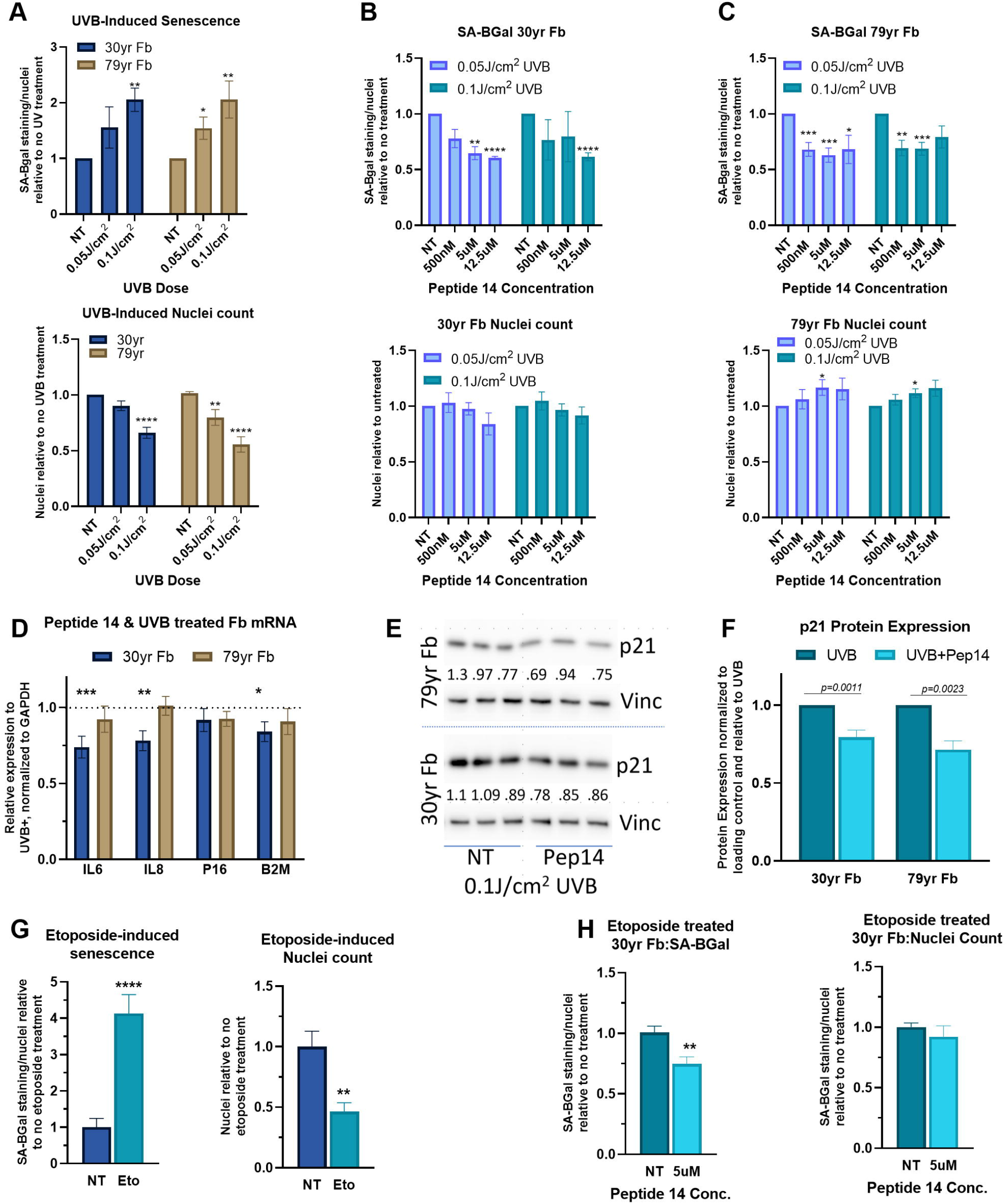
Peptide 14 protects cells from acute UVB and etoposide induced cellular senescence. (A) Human Dermal Fibroblasts (HDFs) isolated from 30 yr and 79 yr healthy donors were exposed to two doses of UVB radiation and stained for SA-Bgal and cell survival, measured according to the number of cellular nuclei. Relative SA-Bgal staining and cell survival of 30 yr (B) and 79 yr (C) HDFs treated with different concentrations of peptide 14 following UVB exposure. (D) mRNA expression of HDFs treated with 500 nM peptide 14 following 0.1 J/cm2 UVB exposure. Data were normalized to the mRNA expression of untreated HDFs exposed to 0.1 J/cm2 UVB. (E) Representative Western Blot analysis of P21 protein expression in 30 yr and 79 yr HDFs treated with or without 500 nM peptide 14 following 0.1 J/cm2 UVB exposure with Fiji quantification of P21 relative to Vinculin and no treatment. (F) Quantitative analysis of P21 protein expression in 30 yr (four independent experiments) and 79 yr (three independent experiments) HDFs treated with or without 500nM peptide 14 following 0.1 J/cm2 UVB exposure. (G) Relative SA-BGal and cell count of HDFs incubated with 20 μM Etoposide to no treatment. (H) SA-BGal and cell count of HDFs incubated with 20 μM etoposide and treated with peptide 14 at 5 μM, relative to no peptide treatment. Data representative of ≥3 independent experiments, *p<0.05; **p<0.01; ***p<0.001; ****p<0.0001, compared to untreated control, according to student’s t-test.

The genotoxic agent Etoposide was also utilized to induce cellular senescence and further confirm the efficacy of peptide 14. The obtained results confirmed that Etoposide effectively induced cellular senescence and cell death (Fig. 2G), and treatment with peptide 14 significantly reduced cellular senescence levels, as measured by SA-BGal (Fig. 2H).

### Peptide 14 reduces biological age in skin models

The role of cellular senescence in local and systemic aging has been previously described and recognized in the skin^7,9,31^ In order to assess whether peptide 14 treatment would effectively decrease cellular senescence in more complex and representative skin samples, we replicated skin aging *in vitro* by building 3D skin equivalents using fibroblast and keratinocyte cultures derived from elderly donors. The aged phenotype was confirmed by measuring the overall structure and quality of the skin, which decreased according to the donor age (Sup. Fig. 2A). Gene expression analysis revealed that, as cell donor age increased, P16 and IL-8 mRNA expression significantly increased, while Ki67 and HAS-2 mRNA expression tended to decrease (Sup. Fig. 2B).

Rapamycin is a long studied molecule affecting mTOR/nutrient signaling and has recently been shown to decrease P16 levels of aging skin^21^, therefore it was chosen as a positive control of senotherapeutic effect in aging skin models. When added to culture media, peptide 14 promoted the maintenance of the overall structure of the 3D skin equivalents created with cells from 48yr old donors, as analyzed by H&E staining, whereas Rapamycin caused a detrimental effect in overall skin structure, including a a thinner and more disorganized epidermis. The effect of both molecules was quantified according to morphological changes analyzed through an unbiased skin score and showed that, while peptide 14 treatment significantly increased the score, Rapamycin treatment led to a decrease (Fig. 3A). The mRNA expression of the peptide 14-treated epidermis from 3D skin equivalents built from donors aged 32, 48, and 60 years exhibited a significant decrease in P16 and a trend towards decreased IL6, in addition to a significant increase in Keratin 1 (a marker of keratinocyte terminal differentiation) and 14 (a marker of non-differentiated, proliferative keratinocytes) (Fig. 3B). Rapamycin induced a significant increase in P16 expression, a trend towards increased expression of inflammatory markers (IL6 and IL8), and a significant decrease in Keratin 1 gene expression levels (Fig. 3B). In the dermis, peptide 14 treatment promoted a significant reduction in B2M gene expression, a pro-aging factor, as well as in the expression ofIL8. Rapamycin treatment induced no significant changes in these markers and increased Matrix Metalloproteinase-1 (MMP1) gene expression, indicative of breakdown of the extracellular matrix (Fig. 3C). Rapamycin and peptide 14 significantly reduced gene expression of Hyaluronidase-1 (HYAL1), which plays a role in the degradation of hyaluronic acid (Fig 3C). Unlike Rapamycin, peptide 14 treatment stimulated increased gene expression of Collagen 1 and Hyaluronan Synthase-2 (HAS2) (Fig. 3C). Therefore, compared to Rapamycin, peptide 14 treatment of 3D human skin equivalents significantly improved numerous markers of skin health and longevity.

**Figure 3.**
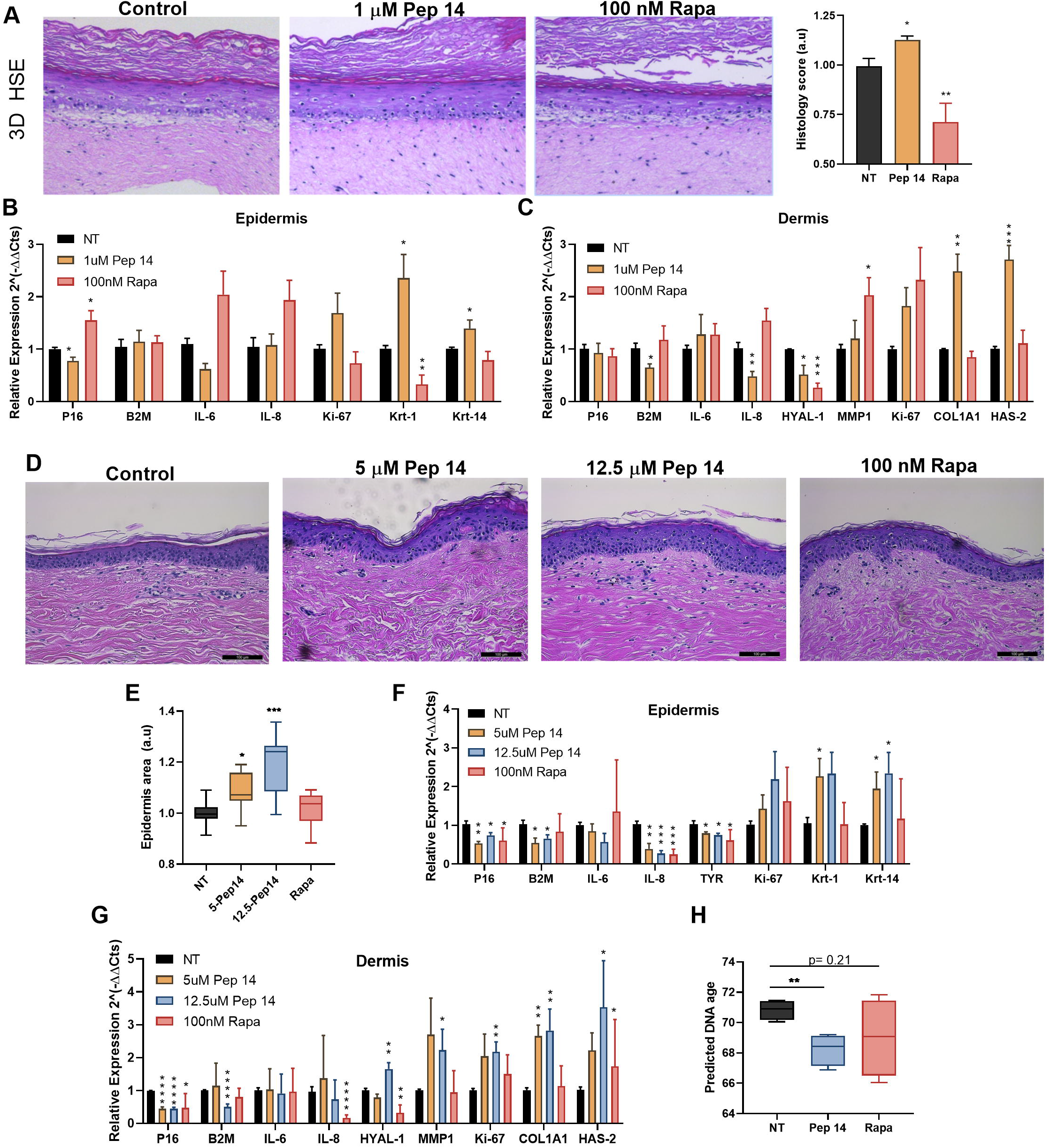
Peptide 14 reduces biological age in skin models. Three-dimensional human skin equivalents (3D HSE) were built using primary cells from chronologically aged donors (32, 48 and 60yr) and treated with basal media, 1 μM peptide (Pep) 14 or 100 nM Rapamycin (Rapa) added in the media twice (day 0 and 3) for 5 days. (A) Representative Hematoxylin and Eosin staining of histological sections of control, peptide 14 and Rapamycin-treated 3D HSE (48y) after 5 days of starting treatment. Histological scores of the 3D HSE, as analyzed by experimental group-blinded personnel (graph on right). mRNA expression of the epidermal (B) and dermal layers (C) of control, peptide 14 and Rapamycin-treated 3D HSE. (D) Representative Hematoxylin and Eosin staining of histological sections of ex vivo skin samples from 55yr donor maintained in basal media, or treated with 100 nM Rapamycin (Rapa), 5 μM peptide 14, or 12.5 μM peptide 14, added in the media. (E) Epidermal thickness analysis of ex vivo skin samples (35, 55 and 79yr) maintained in basal media, or treated with 100 nM Rapamycin (Rapa), 5 μM peptide 14, or 12.5 μM peptide 14. mRNA expression of epidermal (F) and dermal (G) layers of treated samples (35, 55 and 79yr). (H) DNA methylation age calculated using the Skin-Specific DNA Methylation Clock MolClock of ex vivo skin samples (79yr) maintained in basal media, or treated with 100 nM Rapamycin (Rapa), or 12.5 μM peptide 14, added in the media. Data representative of ≈3 independent experiments, *p<0.05; **p<0.01; ***p<0.001; ****p<0.0001, compared to untreated control, according to student’s t-test.

To support the results observed with peptide 14 treatment in human skin equivalents, we treated *ex vivo* skins from donors aged 35, 55, and 79 years. Histological imaging depicted a significant increase in epidermal area with increasing concentration of peptide 14 and no significant changes were observed with Rapamycin treatment compared to control (Fig. 3D, E). In the epidermis of treated skins, both peptide 14 and Rapamycin decreased P16 mRNA expression yet only peptide 14 treatments decreased B2M expression (Fig. 3F). IL8 and tyrosinase (TYR) gene expression were significantly reduced in all treatments though an increase in Keratin 1 and 14 was only observed with peptide 14 treatment (Fig. 3F). In the dermis, peptide 14 treatment more drastically decreased P16 mRNA expression compared to Rapamycin treatment and only the higher concentration of peptide 14 was able to reduce B2M gene expression (Fig. 3G). Rapamycin treatment did reduce both IL8 and HYAL1 gene expression and, similarly to peptide 14, was able to increase HAS2 expression (Fig. 3G). Peptide 14 treatment also increased dermal Collagen 1 and the proliferation marker Ki67 in dermis though did increase HYAL1 and MMP1 gene expression in the higher concentration (Fig. 3G).

In order to have a more in depth understanding of how both peptide 14 and Rapamycin treatments might alter skin’s biological age, the 79 year old *ex vivo* skin biopsies were processed for DNA isolation and methylome analysis. Using the Skin-Specific DNA methylation algorithm developed by our group^32^, we observed that while Rapamycin treatment did not generate any significant alterations (p=0.21), peptide 14 treatment significantly reduced the DNA methylation age (p<0.01) (Fig. 3H).

### Peptide 14 acts as a senotherapeutic agent by regulating aging-related pathways

Our *in vitro* and *ex vivo* data determined that peptide 14 acts as an effective senotherapeutic agent to reduce cellular senescence and biological age. To understand the underlying mechanism of action of the peptide, we incubated fibroblasts isolated from healthy and HGPS donors with peptide 14 for 24 and 48 hours, respectively, and processed the samples for RNA-Seq analysis. We obtained an average of 70 million reads per sample. More than 1.2 billion good quality reads (corresponding to more than 95% of bases showing quality superior to Q30 in the Phred scale, with mean quality of Q36) were generated for all libraries (Supplementary Table 1). The majority of the reads (67%) were uniquely mapped with high quality to the human genome (Supplementary Figure 3A,B). Since the effect on gene expression were more pronounced on HGPS fibroblasts probably due to the longer stimuli, we used this dataset as our gold standard regarding the gene signature modulated by the treatment with peptide 14 (Supplementary Figure 3C and D).

After normalization, no differentially expressed genes were detected among conditions under FDR < 0.05 after correction for multiple-hypothesis. We therefore checked differences in basal gene expression of the top 20 genes (p-value < 0.05, Supplementary Table 2) that were modulated in HGPS fibroblasts by peptide 14 treatment (Fig. 4A). We then compared the gene expression patterns of HGPS fibroblasts with fibroblasts derived from a 41 year-old donor after peptide treatment and saw similar, though not as striking, gene expression signatures (Fig. 4B).

**Figure 4.**
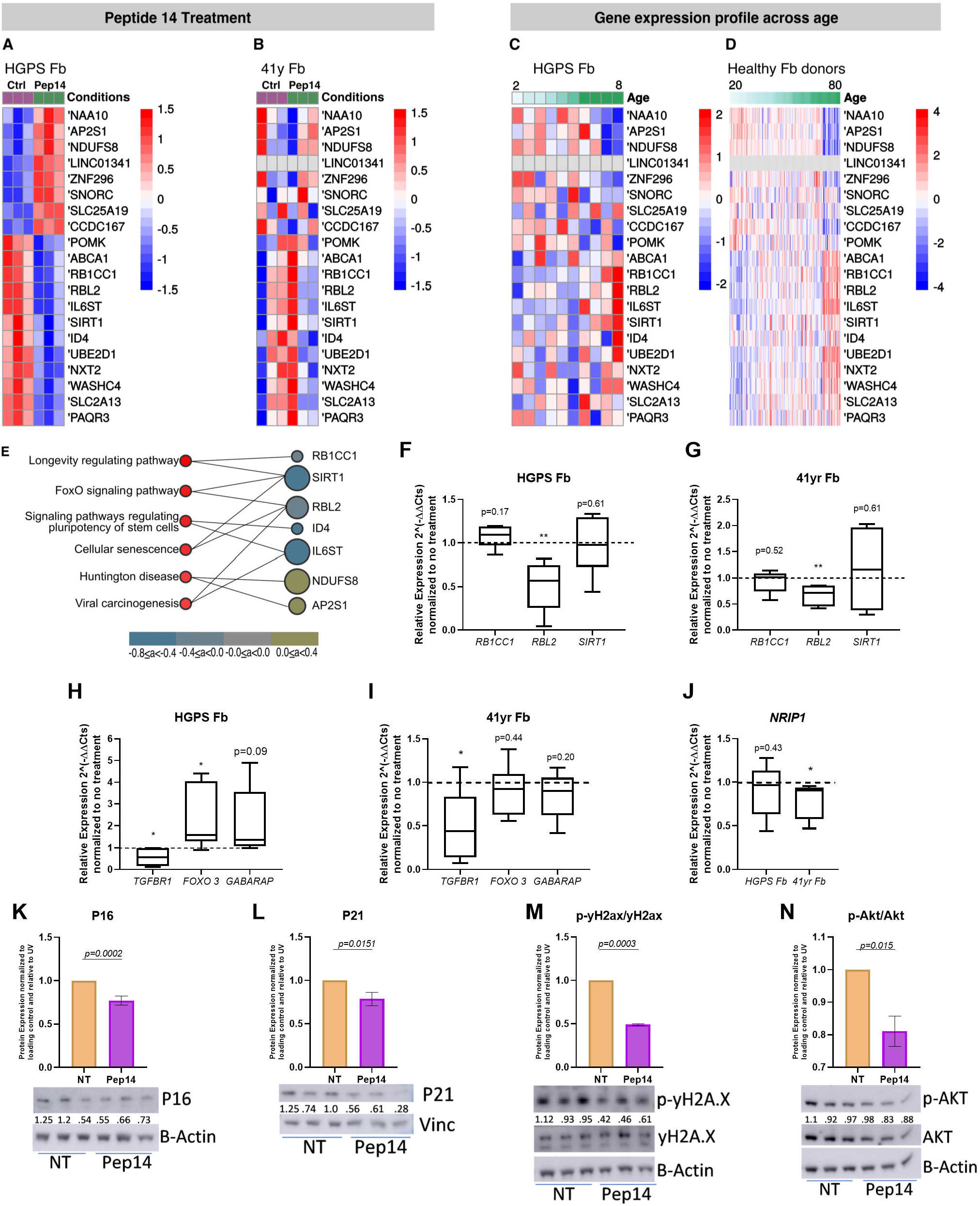
Peptide 14 acts as a senotherapeutic agent by regulating aging-related pathways. Heat maps showing the expression pattern of top 20 genes among different experimental conditions in HGPS (A) and 41yr (B) HDFs. For the comparison between peptide 14-treated (12.5 uM) and control groups, samples were hierarchically clustered using distance as 1 - Pearson correlation coefficient. Heat maps of HDFs samples derived from HGPS patient (C) and healthy donors (D) sorted according to the donor’s age. Color codes in all heat maps represent RNA-seq normalized pseudocounts in log2 scale after row-wise z-score transformation. (E) Pathway Enrichment analysis view associated with some of the top 20 genes modulated by peptide 14 treatment. Only pathways with p-value < 0.05 were represented. Genes are represented by circles of different sizes, reflecting the number of pathways they are involved with. Genes were also colored from blue to yellow according to their log fold change (LFC) in samples treated with peptide 14 compared to untreated controls. Image was modified from Pinet server. qRT-PCR analysis of some of the top 20 genes modulated by peptide 14 treatment in HDFs obtained from HGPS(F) and 41yr HDFs (G) treated with 12.5 μM peptide 14. qRT-PCR analysis of some of the top 89 genes modulated by peptide 14 treatment in HDFs obtained from HGPS (H) and 41yr HDFs (I) treated with 12.5 μM peptide 14. (J) NRIP1 mRNA expression in HGPS and 41yr HDFs treated with 12.5 μM Peptide 14. Protein expression analysis of P16 (K), P21 (L), pyH2ax/yH2ax (M), and pAkt/Akt (N) in HDFs obtained from HGPS HDFs and treated with 12.5 uM peptide 14. Data representative of =3 independent experiments, *p<0.05; **p<0.01, compared to untreated control, according to student’s t-test.

To investigate whether the genes modulated by peptide 14 were associated with aging, we used publicly available HDF RNA-seq datasets of ten samples obtained from HGPS patients and 133 samples derived from healthy donors aged between 1 and 94 years^33^. The heat maps showed that, with few exceptions, the top 20 genes modulated by peptide 14 reflected similar expression pattern changes during aging, and that peptide 14 treatment promoted a “rejuvenated” gene expression signature in treated samples (Fig. 4C,D). Interestingly, gene enrichment analysis of the top 20 genes modulated by the peptide treatment displayed pathways associated with cellular senescence, longevity, and FoxO signaling, among others (Fig. 4E and Supplementary Table 3).

To have a broader overview, a larger signature consisting of the top 89 genes modulated by peptide 14 (p value < 0.1) in HGPS fibroblasts were also evaluated and compared to age-related gene expression changes. Most of the genes in the extended signature were also shown to be modulated in 41yr HDFs upon peptide 14 treatment (Supplementary Fig. 4A,B). Similarly to what was observed in the top 20 genes, the top 89 genes reflected similar expression pattern changes during aging (Supplementary Fig. 4C,D). Pathway Enrichment analysis using the extended signature identified additional pathways including endocytosis, TGF-beta signaling and Th17 cell differentiation, besides FoxO signaling (Supplementary Table 4). Interestingly, the genes associated with the Cellular Senescence pathway, *RBL2 (*LogFoldChange - LFC=-0.3, p value=0.04), *SIRT1 (*LFC=-0.43, p value=0.04), and *TGFBR1 (*LFC=-0.52, p value=0.05) also take part in the FoxO signaling, highlighting this as an important pathway regulated by peptide 14. In fact, peptide 14 was able to modulate several genes associated with both Cellular Senescence and FoxO Signaling pathways annotated according to the Kyoto Encyclopedia of Genes and Genomes (KEGG) database. Since FoxO Signaling is also associated with Rapamycin treatment, we sought to check whether the Rapamycin treatment in both HGPS and healthy donor-derived HDF samples modulated the gene expression in the same direction as the peptide 14. Only three genes presented similar mRNA expression alterations (Supplementary Figure 4E,F). This result suggests that, although regulating some pathways in common, the mechanism of action of peptide 14 and Rapamycin are unique.

To further investigate the molecular mechanism of peptide 14, we searched for consensus small molecule gene expression signatures that mimicked the extended peptide 14 treatment signature. The top two small molecules that displayed the highest number of overlapping perturbed genes were YM-155 and Curcumin (Supplementary Fig. 4G and Supplementary Table 5), previously shown to reduce senescence^34,35^.

Real-time qPCR was utilized to validate the top hits from the RNA-seq results and observed a significant reduction in *RBL2* gene expression with peptide 14 treatment in both HGPS and 41yr HDFs though no statistical significance was observed in *RB1CC1* or *SIRT1* (Fig 4F,G). Peptide 14 treatment significantly decreased *TGFBR1* gene expression in both HGPS and 41yr HDFs though *FOXO3* expression was only upregulated in HGPS HDFs (Fig. 4H,I). *NRIP1* gene expression was validated and, in response to peptide 14 treatment, a significant reduction was observed in 41yr HDFs (Fig. 4J).

We then measured alterations in protein levels of key senescence markers, P16 and P21, and observed that peptide 14 treatment significantly reduced protein markers in HGPS HDFs (Fig. 4K,L). Upstream of P16 and P21, the phosphorylation at Ser139 of yH2AX is an early marker of DNA damage and double stranded DNA breaks. Corroborating the P16 and P21 data, peptide 14 induced a significant decrease in the phosphorylation of yH2AX in HGPS HDFs (Fig. 4M). Lastly, as our RNA-seq analysis highlighted the influence of peptide 14 on FoxO, longevity, and cellular senescence pathways, we investigated its potential to modulate Akt, a member of the PI3K/Akt/mTOR pathway that integrates nutrient, stress, and energy signals to control cell growth, proliferation, and metabolism, among others^36^. This pathway, as well as that of p53/p21 and Bcl-2/Bcl-XL, were recently identified as pro-survival pathways that, when inhibited, result in a reduction of senescence in mouse and human cells^37^. In HGPS HDFs, peptide 14 significantly reduced the phosphorylation of Ser473 of Akt (Fig. 4N), providing indications for further elucidation of the mechanism of action of peptide 14. Combined, these results highlight peptide 14 as a senotherapeutic agent, and that its mechanism of action is associated with AKT/FoxO signaling and pathways involved in senescence and longevity.

### Topical application of peptide 14 in ex vivo skins decreases cellular senescence and strengthens skin barrier

Senescent cell accumulation in the skin has been related to facial wrinkling and perceived age^31^. Since peptide 14 was observed to reduce cellular senescence in human skin models when added to the culture media, we investigated whether a topical peptide 14 treatment would also result in improved skin structure and molecular health. We created a formulation that supported the penetration of up to 2% of the peptide into the dermis (data not shown). Therefore, the concentration of peptide 14 in the formulation was adjusted accordingly (0.01% w/v). Here, topical Retinol was used as an “anti-aging” control, due to its widespread and decades-long use in the field^38–40^.

The treated *ex vivo* skin biopsies (from donors 35, 55, and 79 years of age) were analyzed according to the overall structure of the skin and gene expression of key aging and skin health related markers. Interestingly, topical peptide 14 promoted a significant increase in epidermal thickness (Fig. 5A), a parameter which is associated with the improvement in the appearance of wrinkles^41–43^ and a stronger skin barrier^44^. This contributes to the prevention of water loss in the skin and even improvement of organismal health^45^. Gene expression of the topical peptide 14-treated *ex vivo* samples revealed a significant decrease of P16 mRNA expression in the epidermis of the skins, while topical retinol produced significant increase in both the epidermis and dermis. B2M expression was also significantly decreased with topical peptide 14 treatment in epidermis and dermis, and only in the dermis after Retinol treatment. In the epidermis, topical peptide 14 significantly reduced TYR expression and did not alter inflammation levels of IL6 and IL8, while Retinol significantly increased IL8. The proliferation marker Ki-67 was significantly increased in the epidermis in both treatments, but Retinol significantly reduced both keratin 1 and 14 (Fig. 5B). Topical peptide 14 and Retinol treatments significantly induced dermal Collagen 1 and the Ki67 expression, but only topical peptide 14 treatment induced a significant reduction in the expression of HYAL-1. An increase in HAS2 was only observed in the Retinol treated dermal samples (Fig 5B).

**Figure 5.**
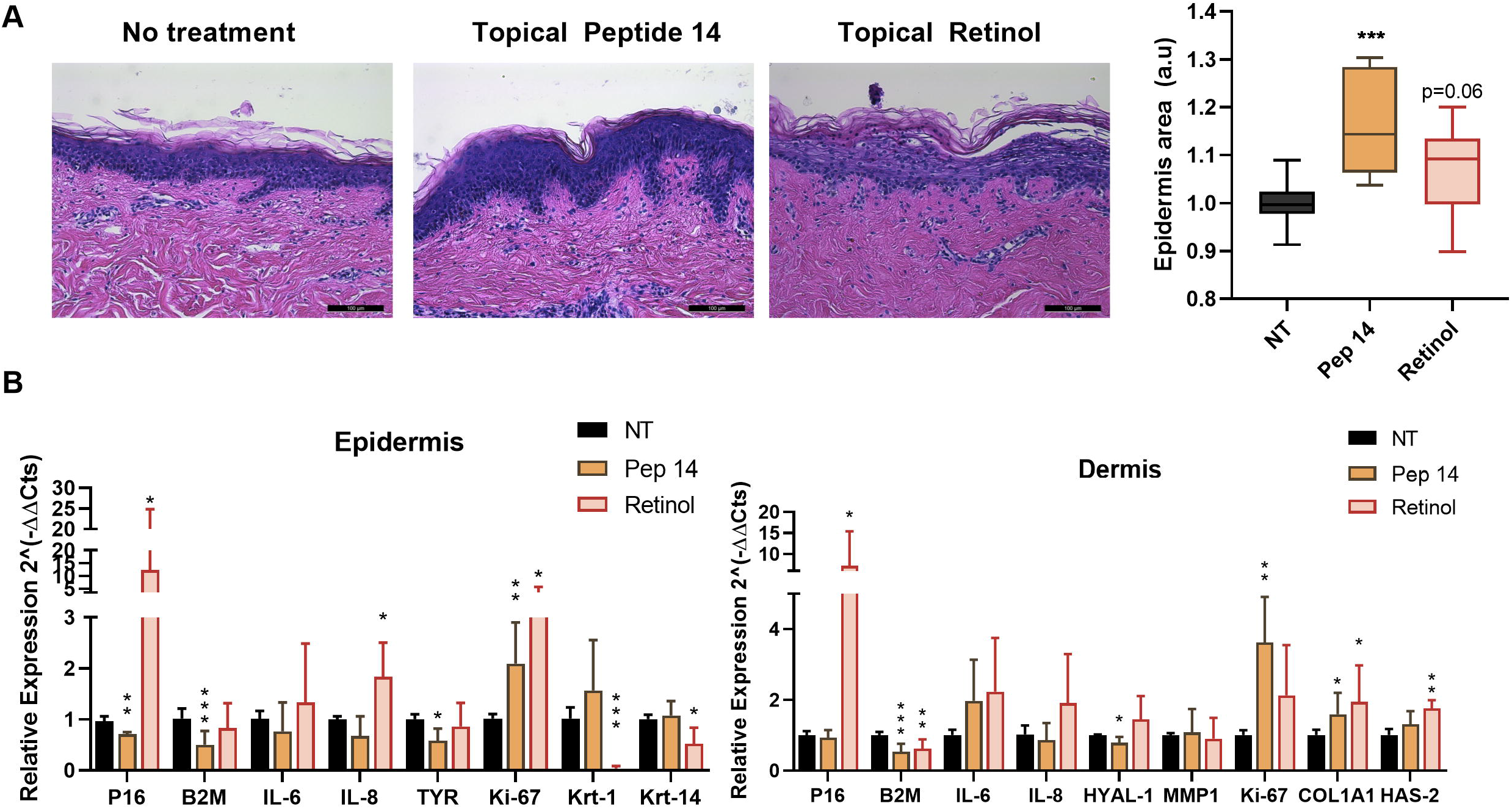
Topical application of peptide 14 in ex vivo skin samples decreases cellular senescence and strengthens skin barrier. (A) Representative Hematoxylin and Eosin staining of histological sections of ex vivo skin samples from 35yr donor maintained in basal media, or treated with topical peptide 14 or topical Retinol. (B) Epidermal thickness analysis of ex vivo skin samples (35, 55 and 79yr) maintained in basal media, or treated with topical peptide 14 or topical Retinol. mRNA expression of epidermal (C) and dermal (D) layers of treated samples (35, 55 and 79yr). Data representative of ≥3 independent experiments, *p<0.05; **p<0.01; ***p<0.001; ****p<0.0001, compared to untreated control, according to student’s t-test.

### *Peptide 14 is safe for long-term use, increases median and maximum lifespan, and improves healthspan in* Caenorhabditis elegans *worms*

Since the senotherapeutic effect of peptide 14 was confirmed in cells, 3D skin equivalents, and *ex vivo* skin samples, we assessed whether the peptide had any mutagenic properties. No changes in toxicity, carcinogenesis, or mutagenicity was observed by Ames test, micronucleus assay, or karyotyping analysis (data not shown). We then assessed whether peptide 14 was safe for long-term treatment using a classic model organism of longevity, *Caenorhabditis elegans* (*C.elegans*)^46–48^. Even though *C. elegans* have been used for decades for toxicological studies and to identify potential candidates for lifespan and healthspan extension, maximum lifespan of the N2 strain of worms often varies from lab to lab due to geographic location, handling, temperature, and food^49,50^. Therefore, two independent research groups ran independent analyses to investigate how peptide 14 alters survival and activity of *C. elegans*. The Site 1 research group is located at Unicamp in Brazil and the Site 2 research group is located at NemaLife in Texas. Lifespan and healthspan were monitored comparing peptide 14 treatment, added to the media, to a vehicle control using both the standard agar/bacteria plate-requiring routine manipulations of worms by technicians (Site 1), and a microfluidic chamber mimicking a soil-like environment and computer scanning for movement analysis (Site 2). Regardless of the study location or methods utilized, peptide 14 did not pose any toxic effects to the worms and were able to significantly increase median worm lifespan by 15% at Site 1 and 12.5% at Site 2 and extended overall lifespan compared to controls by an average of 2 days at both sites (Fig. 6A,B). A 1 μM peptide 14 concentration led to the best results at both sites, though both 0.1 and 10 μM also resulted in increased lifespan (data not shown). In addition to the benefits of lifespan extension observed in worms treated with peptide 14, healthspan measures also improved. At Site 1, 1 μM peptide 14 treatment significantly increased movement of worms by measuring thrashing per minute in liquid media (Fig. 6C). Site 2 also observed that peptide 14-treated worms had a higher percentage of highly active individuals, compared to the vehicle control (Fig. 6D), and that peptide 14-treatment induced a shift in the population of active worms, compared to control. Even by day 12, peptide 14-treated worms were more active than controls at day 5 (Fig. 6E).

**Figure 6.**
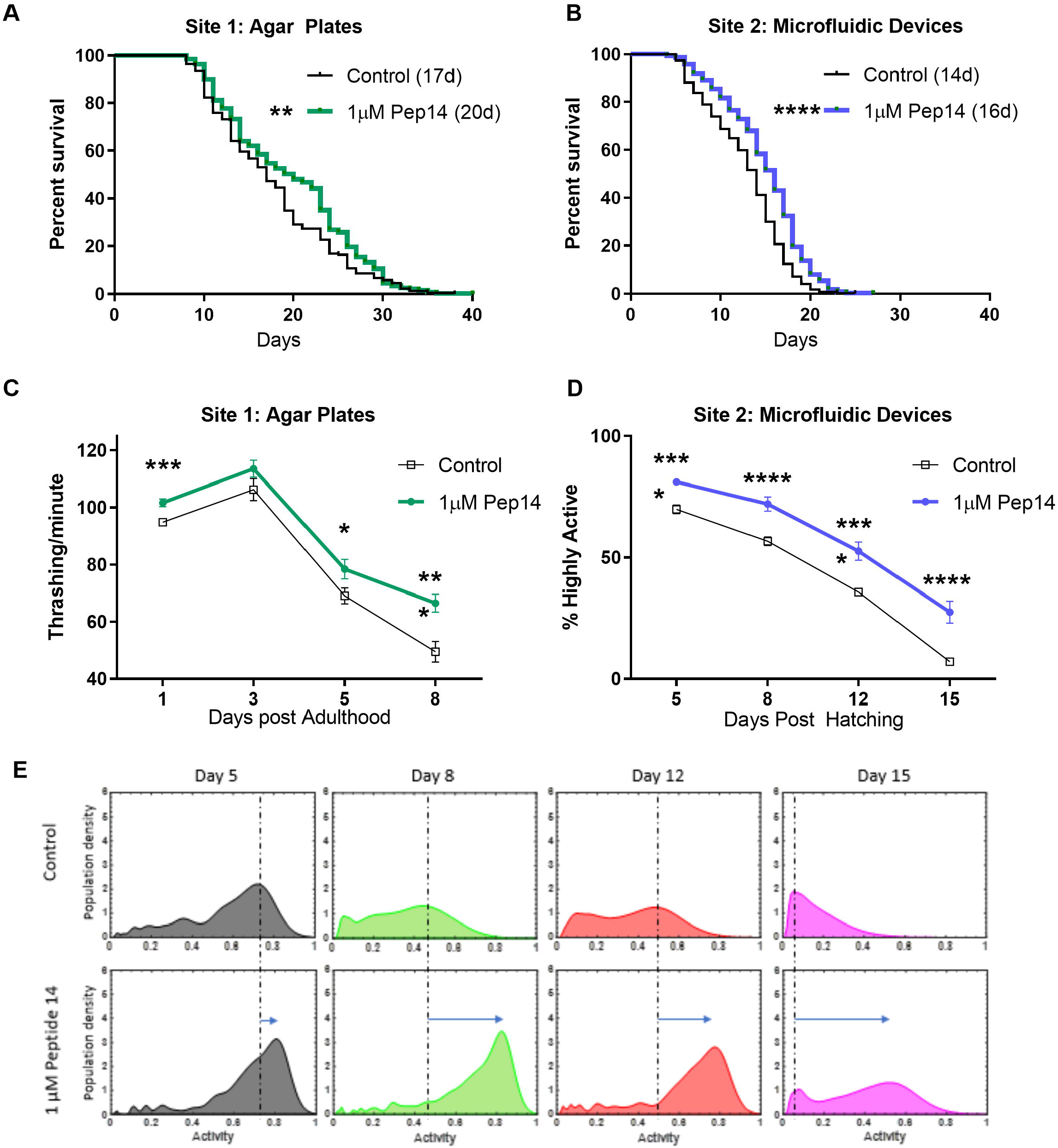
Peptide 14 increases median and maximum lifespan and improves healthspan of Caenorhabditis elegans worms. (A) Lifespan analysis of control and peptide 14-treated worms grown in agar plates. Median lifespan is presented within parenthesis next to group identification. Each line represents the mean of 3 independent pooled experimental replicates of 120 worms per experimental group. (B) Lifespan analysis of control and peptide 14-treated worms grown in microfluidic devices. Median lifespan is presented within parenthesis next to group identification. Each line represents 3 independent experiments of approximately 150 worms per experimental group. (C) Manual analysis of worm movement in liquid media (Thrashing). Thrashing was measured on 45 worms per experimental group, which came from 3 independent experiments. (D) Percentage of highly active worms grown in microfluidic devices. Data refers to 3 independent replicates of 130-150 worms per experimental group. (E) Comparative analysis of the mobility of untreated (control) and peptide 14-treated worms grown in microfluidic devices in different experimental time-points. Dashed line indicates peak worm activity levels in the control group and arrows indicate the shift in highly active worms with peptide 14 treatment, *p<0.05; **p<0.01; ***p<0.001; ****p<0.0001, compared to untreated control, according to the Log-rank test.

## Discussion

Aging is associated with compromised tissue function and loss of resilience and regeneration upon damage. In the skin, senescent cells accumulate due to intrinsic and extrinsic factors^16^. While intrinsic skin aging is associated with epidermal and dermal thinning, loss of epidermal rete ridges (which decreases the interface area of epidermis and dermis), and vascularization^16^, extrinsic skin aging is associated with aggravated extracellular matrix loss, and the formation of abnormally elastic tissues^16^. Both are characterized by senescent cell accumulation. Senescence-inducing intrinsic factors include time (i.e. chronological aging)^5–8^, and genetic background, as observed in patients with genetic diseases such as Werner syndrome^51^, and Hutchinson-Gilford Progeria Syndrome^52^. Senescence-inducing extrinsic factors include UV radiation^26,27^, smoking^53^, and pollution^54^, to name a few. When chronically accumulated in the skin, such cells compromise epidermal cell renewal^18^, skin regeneration^55^, and ECM homeostasis^56^ correlating with the perceived age of the skin^7,31^. Measuring the approximate number of senescent cells in the skin can be challenging and current studies indicated that epigenetic changes comprise a significant component of the aging process. Therefore, we proposed the use of a Skin-Specific DNA methylation clock that can offer a comprehensive and highly accurate alternative method to analyze human skin health status and aging^32^.

Few recently published studies have demonstrated that reducing senescence burden improves skin function. Methylene blue treatment decreased aging markers, including SA-Bgal and P16 expression in human fibroblasts treated for 4 weeks. The treatment of 3D skins with methylene blue increased the skin viability, dermal thickness, and extracellular matrix protein expression^57^. Another study using 3D skin models identified an alcoholic extract of *Solidago alpestris* with the ability to block the negative effects of senescence in human skin fibroblasts including SASP and improved epidermal stratification. Although those reports reinforce the benefit of targeting senescence to improve skin function, the molecules can present high toxicity and challenges to be obtained in large scale and consistent composition. Peptides are multifunctional, pleiotropic molecules that are known to be non-toxic^58^. Within and between cells, peptides frequently act as signaling molecules, participating in numerous physiological functions. Therefore, such class of biomolecules has long been seen as an alternative therapeutic intervention able to mimic natural pathways^59^. In the present study, we implemented a screening platform based on human skin cells and tissues, which faithfully recapitulate skin aging *in vitro*, therefore increasing the chances of finding molecules with translational benefit to human skin.

After a comprehensive and optimized screening process, peptide 14 was shown to significantly reduce cellular senescence in different senescence-inducing experimental models, which recapitulate both intrinsic (e.g. chronologically aged and HGPS cells) and extrinsic (UVB radiation and etoposide treatment) cellular senescence stimuli. Since the cellular senescence burden of the skin correlates with tissue structure and function, the treatment of 3D skin equivalents and *ex vivo* skin biopsies with the peptide caused significant reduction in the gene expression of senescence (e.g. P16), inflammation (IL-8), and pigmentation (TYR) markers, as well as an increase in cellular renewal (Ki67 and KRT-14), and terminal differentiation of keratinocytes (KRT-1) markers. Such molecular signatures were accompanied by an increase in health parameters of 3D skin equivalents, including an improvement on the morphological tissue organization which resembles a younger skin profile. Even more striking, the 5-day treatment of *ex vivo* skin samples with peptide 14, induced a significant reduction of an average 2.6 years in the biological age of the tissues. Here, Rapamycin, a well known senotherapeutic^60,61^, was utilized as a positive control. In accordance with previously published studies^21^, the treatment of *ex vivo* skin biopsies with Rapamycin reduced P16 mRNA expression in both the epidermis and dermis, as well as IL-8. Nevertheless, the skin phenotype (i.e. epidermal area) and biological age (DNA methylation age), were either worsened or unchanged. These data reflect that even though Rapamycin is a well-published senotherapeutic molecule recently tested in a topical formulation for reducing cellular senescence in skin^21^, peptide 14 outperforms Rapamycin in both 3D human skin equivalents and *ex vivo* donor skin biopsies according to histological, gene expression, and DNA methylation parameters, reflective of a senotherapeutic molecule that significantly improves skin health and reduces tissue biological age.

Although the mechanism of action in which peptide 14 reduces cellular senescence burden is complex with the regulation of multiple pathways, RNA-Seq suggests that peptide 14 modulates important genes from Akt/FoxO signaling and other genes involved in senescence and longevity pathways. Down-regulated genes included *RB1CC1*, a protein-coding gene involved in cell growth and proliferation, apoptosis and cell migration^62^; and *RBL2* (RB Transcriptional Corepressor Like 2), which codes for a protein that controls gene expression by binding to the E2F transcription factors and recruiting epigenetic modifiers, and is a direct substrate of Akt^63^ *and TGFBR1*, a receptor that activates the transforming growth factor beta (TGF-beta) pathway, that exhibits multifaceted crosstalk with aging processes. TGFBR1 inhibitors were capable of reversing not only the proliferation arrest of senescence induced by the integrin Beta3, but also the upregulation of p21^64^. We observed increased *SIRT1* gene expression with age in HDFs, as has been previously described in aged or acutely damaged tissues^65^, and peptide 14 treatment reduced SIRT1 gene expression. We also compared the RNA signatures of fibroblasts treated with peptide 14 and rapamycin. Although we observed that both treatments regulate some common pathways such as FoxO signaling, the mechanism of action of peptide 14 and Rapamycin are unique.

Our data suggests that peptide 14 reduces the accumulation of DNA damage and prevents cellular senescence via SASP, acting through FoxO and cellular senescence signaling. These common pathways are shared amongst other senotherapeutic molecules such as Rapamycin, Curcumin, and YM-155. YM-155 is also known as Sepantronium bromide, and is a survivin inhibitor, while Curcumin, also known as Diferuloylmethane, is a natural phenolic compound and a p300/CREB-binding protein-specific inhibitor of acetyltransferase. The gene *NRIP1* (Nuclear Receptor Interacting Protein 1) is modulated by both Curcumin and YM-155 and was the most frequently modulated gene of all small molecules analyzed when we looked for drugs that were activating similar pathways. Our data demonstrated a decrease in NRIP1 expression after peptide 14 treatment. Interestingly, the deletion of this gene in old mice resulted in increased autophagy, reduced the number of senescent cells and extended female longevity^66^.

The translational application of peptide 14 to the skin was assessed by investigating the effect of a topical formulation in *ex vivo* skins. The topical peptide 14 treatment was compared to Retinol, since retinoids are considered reference “anti-aging” active ingredients in the cosmetic industry, due to the accumulated knowledge regarding their effects over mammalian cells and skin^2,38–40,67–70^. Strikingly, in our hands, Retinol treatment of *ex vivo* skin samples promoted the expected peeling effect of the stratum corneum, but also promoted a significant increase in P16 mRNA expression in both the epidermal and dermal layers. Furthermore, Retinol induced a significant increase in IL8 mRNA expression, which is likely associated with the irritation commonly caused by this ingredient. Such an observation allows one to suggest that, even though such a molecule promotes peeling and visual renewal of the skin, it may increase inflammation and cellular senescence markers in the skin compromising skin health in the long-term. Further experiments *in vitro* and *in vivo* must be performed to investigate the cellular senescence and biological age status of Retinol-treated skin. Alternatively, topical peptide 14 increased epidermal thickness, decreased senescence markers, maintained lower levels of inflammation, and induced cell proliferation and collagen 1A1 expression, leading to a healthier skin profile.

Many cellular senescence and longevity signaling pathways are intertwined and conserved throughout evolution^71^. Corroborating such a notion, peptide 14 treatment of *C. elegans* indicated that the molecule was able to extend the lifespan of treated worms and the effects on healthspan parameters were even more impressive. Since cellular senescence contributes to age-related tissue deterioration of different body sites, it is possible that the observed effects derive from the senotherapeutic potential of the peptide. Nevertheless, additional studies are necessary to elucidate the application and specific mechanisms of action of the peptide in other tissues and in the whole organism.

Taken together, these data demonstrate that peptide 14 is a novel senotherapeutic compound that not only improves skin health, but is safe for long-term use and is able to extend the lifespan and healthspan of *C. elegans*. By using a screening platform based on the cellular senescence phenotype, we showed that it is possible to develop a novel therapeutic peptide that is able to both ameliorate signs of skin aging and act as a putative geroprotector that is potentially capable of improving the overall health of an organism.

## Materials and Methods

### Peptide library construction

A peptide library containing 164 short amino acids peptides from a previous work^22^. The library was synthesized using the SPOT methodology with a functionality of 100nmol (Kinexus, Canada). The peptides were diluted in water overnight, under agitation, and then diluted in culture media to a working solution of 50uM.

### Amino acid scanning

The four lead peptides from the peptide library were subjected to an amino acid scanning, where each position was replaced by each one of proteinogenic amino acid residues, generating variants with a point mutation. The new library consisted of 764 new peptides that were synthesized using the SPOT methodology (Kinexus, Canada).

### Chemical synthesis of peptides

The top hits peptides selected (peptide 14, 144, 156, 195, and 393) from the screening were purchased from CPC Scientific Inc. (USA), which synthesized the peptide by solid phase (Fmoc) on a Rink amide resin, with > 95% purity, at the form of acetate salt. The molecular mass was confirmed by mass spectral analysis and purity by the RP-HPLC chromatogram.

### Cell culture

Primary human dermal fibroblasts derived from HGPS donors were obtained from The Progeria Research Foundation Cell and Tissue bank. Healthy normal human dermal fibroblasts and human keratinocytes were either purchased from Coriell Institute for Medical Research (Camden, NJ), MatTek Life Science (Ashland, MA), or isolated from *ex vivo* human skin explants obtained from ZenBio (Research Triangle, NC).

The cells purchased from Coriell Institute for Medical Research included HDF 71 yr (AG05811, XX, arm, caucasian), HDF 84 yr (AG11725, XX, arm, caucasian) and HDF 90 yr (AG08712, XX, arm, caucasian).

Cells purchased from MatTek Life Science included HDF 60 yr (F13400A, XX, african-american), keratinocytes 60yr (K13400A, XX, african-american), neonatal HDFs (F90800, XY, foreskin, caucasian), neonatal keratinocytes (K90800A, XY,foreskin, caucasian).

All other cells were isolated from *ex vivo* human skin obtained from ZenBio (Research Triangle, NC). All skin samples were from XX donors, caucasian, and abdomen area. The ages included 30, 35, 41, 48, 55 and 79 years. The cell isolation was performed as described by Zonari et al.^72^. Briefly, the tissue samples were cut into small pieces of 0.5 cm^2^ and incubated in PBS containing dispase (2.5 U/mL, BD Biosciences) overnight at 4°C. The epidermis was then mechanically separated from the dermis and incubated in 0.5% trypsin-EDTA (Gibco, USA) for 7 min at 37°C to isolate the keratinocytes. The cells were separated from the remaining tissue using a 100 mm pore size cell strainer (BD Biosciences, USA) and the cell suspension was centrifuged at 290 g for 5 min. Human keratinocytes were seeded at a density of 100,000 cells/ cm^2^ in Keratinocyte Serum Free Medium (KSFM) supplemented with Epidermal growth factor and Bovine pituitary extract (Gibco). For the isolation of HDF, the dermis separated from the epidermis was incubated in PBS containing collagenase IA (250 U/mL, Sigma) for 3 hr at 37°C. The HDF were separated from the remaining tissue using a 100 mm pore size cell strainer, centrifuged at 300 g for 5 min and seeded at a density of 50,000 cells/cm^2^ in Dulbecco’s Modified Eagle Medium (Invitrogen, Carlsbad, CA),supplemented with 10% v.v. Fetal Bovine Serum (FBS; VWR) and 1% v.v. Penicillin-Streptomycin (Invitrogen).

### 2D screening of senotherapeutic peptides

HGPS HDFs were seeded in 96 well plates at a density of 4 x 10^3^ cells/well at least 6 h prior to treatment. The peptide library was added at a final concentration of 50 μM in DMEM without FBS. After 30 min, 10% FBS f.c. was added. The cells were incubated for 48 hours at 37°C and 5% CO2. After incubation, cells were fixed for SA-Bgal staining. ABT-263 (ApexBio, final concentration: 5 μM) was added as a positive control. Cells without treatment were considered negative control.

### 3D skin model production and treatments

3D skin models were prepared as described by Maria-Engler et al.^73^, with minor modifications. Briefly, collagen I gels containing embedded fibroblasts were seeded with keratinocytes and cultured for 24 hrs. Then, gels were raised to an air-liquid interface and kept for additional 7 days, to allow epidermal cornification. Treatments were performed by adding the molecule in the culture media or within a topical cream. In 3D skin equivalent assays, Peptide 14 was used at 1 μM, and Rapamycin at 100 nM. Treatment was added to the culture media at day 0 and day 3. Topical peptide 14 treatment consists of 0.01% peptide 14 in a 15% oil moisturizing emulsion. After 5 days, the samples were harvested and fixed in formalin for histology, or used for RNA isolation.

### 3D skin model quality control

Histology of produced 3D skin models was performed as quality control. To do so, 3D skin models are fixed overnight in 10% Formalin, then embedded in paraffin, sectioned and stained with hematoxylin and eosin. Quality assessment follows an internal protocol, in which the analysis of several aspects of the obtained models is performed. Seven different parameters including general organization of cell layers, stratification of epidermis, thickness of corneous layer, among others are evaluated by a blinded person and ranked from 0-4. The maximum score is 28.

### Ex-vivo skin samples and treatments

Skin samples from healthy donors (XX, caucasian, ages 35, 55 and 79 years old) were obtained from ZenBio (Research Triangle, NC). Samples were cultured in an air-liquid interface using DMEM supplemented with 10% v.v. FBS and treatments added either in the media or by topical application using sterile swab. Treatment was added to the culture media at day 0 and day 3. Topical treatment (described above) was also added on day 0 and day 3. After 5 days from the first treatment, the samples were harvested and fixed in formalin for histology, or used for RNA and DNA isolation.

In some experiments, dermal and epidermal layers were separated prior to processing by incubating overnight the skins in dispase solution 2.5 U/mL (Invitrogen). The epidermis was mechanically separated from the dermis using forceps.

### Photoaging experiments with UVB

Cultured fibroblasts were washed with 1x PBS and a small layer left on top. Lids were removed and cells dosed with 0.05 or 0.1 J/cm^2^ UVB (Honle UVASPOT 400/T solar source with Honle UV Meter to determine dosage, Honle, Germany) and media with or without peptide treatment added back immediately after dosing. Cells were incubated with treatments for 30 hr then either fixed for SA-Bgal staining, RNA isolated for qPCR analysis, or Protein isolated for Western blotting.

### Western Blotting

Cells were homogenized with a Tris-SDS-EDTA Lysis buffer containing protease and phosphatase inhibitors and beta-mercaptoethanol. Samples were boiled and run on SDS-PAGE for western blotting according to standard procedures. Nitrocellulose membranes were probed for p21 (CST 2947, Cell Signaling Technology), p16 (CST 80772, Cell Signaling Technology), ß-Actin (sc-81178, Santa Cruz Biotechnology), Vinculin (sc-73614, Santa Cruz Biotechnology), p-γH2AX (sc-517348, Santa Cruz Biotechnology), γH2Ax (sc-517336, Santa Cruz Biotechnology), pAkt (CST 9018, Cell Signaling Technology), and Akt (CST 2938, Cell Signaling Technology). Blots were detected with the Azure Biosystems ECL detection kit (AC2103). Quantification of band intensities by densitometry was carried out using Fiji software.

### Accelerated aging induction in 2D cultured cells

To induce accelerated aging of fibroblasts, cells were treated for 24 h with 20 μM etoposide (Cell Signaling)^37^. Two days after etoposide removal, the cells were treated for 48 h 5 μM of peptide 14 and the senescence level was determined by SA β-galactosidase staining and normalized to total cell number.

### Senescence Associated β-galactosidase staining and quantification

SA β-galactosidase staining was performed using the Senescence β-Galactosidase Staining Kit (Cell Signaling), following manufacturer’s instructions. Briefly, cells were washed with 1x PBS and fixed for 10-15 minutes. Then, cells were washed twice with 1x PBS and incubated with β-Galactosidase Staining Solution overnight at 37°C in a dry incubator without CO2. Cells were then washed and stained with 1 μg/mL of Hoechst 33342 (Invitrogen, H1399) for 10 min and observed by 20x magnification (6D High Throughput, Nikon). The blue staining quantified as the mean colour intensity compared to the total number of cells using CellProfiler^™^. Data were presented as SA β-galactosidase staining levels normalized to total cell number.

### ATRX staining and quantification

ATRX was detected by immunofluorescence as described previously^23^. Briefly, the cells were fixed using 4% paraformaldehyde solution for 10 min. Permeabilization was performed for 5 min using 0.1% Triton followed by blocking for 40 min with 0.5% Tween and 1% BSA. The primary antibody against ATRX (Santa Cruz Biotechnology, D5 - sc55584) was diluted 1:2,000 and incubated overnight at 4°C. After 3 washes with PBS, cells were incubated at room temperature for 1 hour with the secondary antibody Goat Anti-Mouse IgG H&L-Alexa Fluor^®^ 488 (Abcam, Cambridge, MA, ab150113) and Hoechst 33342 (Invitrogen, H1399). Cells were imaged at 40x magnification, using the IN Cell Analyzer 2500 and the IN Cell Developer toolbox (GE Healthcare). The average ATRX foci per cell was defined by the total ATRX foci / total nuclei from a minimum of 150 cells per experimental condition.

### RNA isolation and qRT-PCR

RNA was isolated using RNeasy Mini Kit (Qiagen), following manufacturer’s instructions. Briefly, following different experimental treatments, cells were harvested using 0.25% trypsin-EDTA (Gibco), lysed with lysis buffer and homogenized. Ethanol was added to the lysate and samples were applied to the kit’s spin columns. Contaminants were washed and RNA was eluted in RNase-free water. For qPCR analysis, RNA was quantified using spectrophotometry and 1 μg of RNA of each sample was reverse-transcribed using High-Capacity cDNA Reverse Transcription Kit (Thermo Fisher Scientific), following manufacturer’s instructions. qPCR was performed using PerfeCTa^®^ qPCR ToughMix^®^, Low ROX^™^, (QuantaBio) with Taqman (Invitrogen) probes for CDKN2 (P16) (Hs00923894_m1), B2M (Hs00187842_m1), P21 (Hs00355782_m1), IL-8 (Hs00174103_m1), IL-6 (hS00174131_m1), Ki67 (Hs04260396_g1), Keratin 1 (Hs00196158_m1), Keratin 14 (Hs00265033_m1), TYR (Hs00165976_m1) HAS-2 (Hs00193435_m1), COL1A1 (Hs00164004_m1), HYAL-1 (Hs00201046_m1), MMP-1 (Hs00899658_m1), GABARAP (Hs00925899_g1), FOXO3 (Hs00818121_m1), TGFBR1 (Hs00610320_m1), NRIP1 (Hs00942766_s1), RB1CC1 (Hs01089002_m1), SIRT-1 (Hs01009006_m1), and GAPDH (Hs02758991_g1).

### DNA sample acquisition and methylation analysis

Total DNA samples were obtained from human skin biopsy samples (XX, caucasian, 79 yr) using the QIAamp DNA Mini Kit (Qiagen) and applied to the human Illumina Infinium EPIC 850K chip, as described previously^32^. DNA samples included control, 5 days treatment with 12.5 μM Peptide 14, and 5 days treatment with 100 nM Rapamycin.

### Skin Specific Molecular Clock analysis

Raw methylation files (idat) were converted to beta values using the “minfi” package version 1.32.0^74^ after the quality control and quantile normalized using the function betaqn from the “wateRmelon” package version 1.30.0^75^. Quantile-normalized beta values for all samples were used as input for the Skin-Specific DNAm predictor MolClock^32^. Box plots were used to compare age predictions among groups in different datasets.

### RNA-Sequencing samples preparation

Total RNA samples were evaluated for integrity by using an Agilent 2100 Bioanalyzer (RNA 6000 Nano Chip Total Eukaryotic RNA Assay, RIN ≥9.3) RNA-Seq libraries were constructed using Illumina TruSeq Stranded mRNA Library Prep Kit (Illumina) and sequenced on the Illumina NovaSeq6000 platform in the 100 nt, paired-end configuration.

### RNA-Sequencing analysis

For RNA-Seq analysis, both internal and publicly available data sets were used. SRA files from the project SRP144355^33^ were downloaded and converted to fastq files using SRA Toolkit version 2.8.2-1, while internal data was delivered as fastq files. Reads from raw RNA-Seq data (fastq files) were trimmed using the software Trimmomatic version 0.37^76^ with default options and mapped to the human genome (GRCh38 - ENSEMBL release 88) using STAR version 2.5.3a^77^ with default parameters for single unstranded reads as per developer’s manual. Htseq-count version 0.11.1^78^ was used to assign uniquely mapped reads to genes (excluding pseudogenes) according to the annotation in the Homo_sapiens.GRCh38.89.gtf data set (ENSEMBL release 88). Only genes with a minimum mean of 10 mapped reads were considered for further analysis. Read counts were analyzed using the R package DESeq2 version 1.26.0^79^ and libraries were normalized using the estimateSizeFactors function of the package according to the condition evaluated. For fibroblasts, we evaluated the conditions peptide treatment versus control. Heat maps were constructed using the pheatmap package version 1.0.12 using a regularized Log2-transformed counts-per-million, z-scaled across samples. Genes (rows) were clustered using 1-Pearson correlation coefficient as distance and samples were ordered based on their chronological age, with exception of samples of progeroid fibroblasts (control and treated with peptide 14), that were hierarchically clustered, likewise the genes.

### Pathway enrichment analysis

Gene lists were further projected onto biological pathways of known biological functions or processes enriched using piNET, which holds a comprehensive library of biological gene/protein sets through Enrichr. The Kyoto Encyclopedia of Genes and Genomes (KEGG) 2019 database was used. P-values were controlled for False Discovery Rate (FDR) using the Bonferroni method.

### Gene perturbation signature

To investigate the mechanism of action of peptide 14, we searched for consensus small molecule signatures that mimicked the extended signature of genes modulated upon treatment. The underlying dataset for the search engine included the top 89 modulated genes upon peptide 14 treatment that were compared to a portion of the LINCS L1000 small molecule expression profiles generated at the Broad Institute by the Connectivity Map team (https://maayanlab.cloud/L1000CDS2/#/index/5f5fb98a77dff10054d75ccc). Signatures were scored according to the overlap between the input genes signature and the signature genes from the database divided by the effective input, which is the length of the intersection between the input genes and the L1000 genes.

### C. elegans experiment: Site 1

#### Lifespan analysis

120 wild-type Bristol (N2) worms per replicate were transferred at the first day of their adult life to NGM plates containing 1 μM peptide 14 or controls containing OP50 bacteria and supplemented with 50 μg/mL of the mitosis inhibitor FUdR (floxuridine) to avoid egg-laying/hatching^80^. This experiment was repeated 3 times and by 2 different individuals. Statistical difference was detected by the Log-Rank (Mantel-Cox) survival test, and all groups were compared to the control, no peptide group.

#### Motility analysis

The swimming movement of *C. elegans*, also called thrashing, was measured in liquid M9 buffer media. 15 worms were counted for 30s each. This experiment was repeated 3 times, by 2 different individuals, employing different populations of worms on the listed days after adulthood. Statistical difference was detected by analyzing each individual time point by One-Way ANOVA and Dunnet’s post hoc, and all groups were compared with the control, no peptide group.

### Chip-based C. elegans experiment: Site 2

Unlike the study at Site 1 where lifespan assays were conducted on agar-plates with progeny-blocking drug FUdR, lifespan assays at Site 2 were conducted in microfluidic chips without the use of FUdR (Infinity Chips, NemaLife Inc., TX)^81^.

Wild-type Bristol (N2) C. elegans (n=1811) were used in this study (Caenorhabditis Genetics Center, MN). Peptide 14 at 1 μM was formulated in liquid nematode growth media (NGM) and the food concentration was maintained at 20 mg/mL of E. coli OP50. Three biological replicates of lifespan assays were conducted with each biological replicate consisting of 2 technical replicates. Each technical replica corresponds to a population of > 60 animals in the microfluidic chip. Both survival and motility was scored for the same crawling population in the chip.

#### Lifespan and motility

The life-long assay was initiated by loading Day 1 adults into the microfluidic chips. Subsequent to loading, fresh 1 μM Peptide 14 was administered daily to the worm population in the chips at 20°C, until all worms perished. Videos of crawling animals in the chip were acquired each day prior to feeding fresh Peptide 14 solutions to determine live counts and worm motility. Control worms had no peptide exposure. The videos were analyzed using the Infinity Code software (NemaLife Inc., TX) for worm survival and motility. The number of living worms in the population was determined based on detectable movement. Motility was determined based on the displacement of individual worms from the rectangular area (bounding box) that encloses their whole body. Worms that moved more than their body length within 30 seconds were labeled “highly active.” The percentage of highly active worms in the population was then calculated. Kaplan-Meier curves from the life-long toxicity assay were generated using GraphPad Prism (Version 8.4.2 (679)). Log-rank test was used to compare the survival curves between the non-exposed control and Peptide 14-treated populations.

## Authors’ contributions

Conception and design: AZ, MB, CR, LB, EA, and JC. Peptide library design and lead optimization: OF, WP. Wet lab experiments: AZ, LB, KA, DF, MG, BM. Methylome and RNA seq data analysis: MB. *C. elegans* experiments: MM, MI, WS, MR, TA, SV. Drafting the article: JC, AZ, LB, MB. Final revision: All authors. The authors read and approved the final manuscript.

## Acknowledgements

The authors acknowledge Silvya Stuchi Maria-Engler, Associate Professor at the School of Pharmaceutical Sciences, University of São Paulo, and Paula Comune Pennacchi, post-doc at CNIO, Spain, for their support in the development and optimization of the 3D skin equivalent models.

