## Supplementary material for "Senotherapeutic peptide reduces skin biological age and improves skin health markers": Sup Table 1

Supplementary Table 1 - Statistics and quality control of reads generated for each sample.

| **Sample_ID** | **Condition** | **Cell_type** | **Read Order** | **Index** | **Yield(Bases)** | **Reads** | **% of >= Q30 Bases(PF)** | **Mean Quality Score(PF)** |
| --- | --- | --- | --- | --- | --- | --- | --- | --- |
| R2503_S18_L001 | Control | HGPS - Fibroblasts | R1 | CCGCGGTT-AGCGCTAG | 4,143,733,565 | 41,027,065 | 96.66 | 36.47 |
| R2503_S18_L001 | Control | HGPS - Fibroblasts | R2 | CCGCGGTT-AGCGCTAG | 4,143,733,565 | 41,027,065 | 95.41 | 36.24 |
| R2505_S19_L001 | 48h with 12.5uM Pep14 | HGPS - Fibroblasts | R1 | TTATAACC-GATATCGA | 3,833,050,697 | 37,950,997 | 96.69 | 36.48 |
| R2505_S19_L001 | 48h with 12.5uM Pep14 | HGPS - Fibroblasts | R2 | TTATAACC-GATATCGA | 3,833,050,697 | 37,950,997 | 95.49 | 36.26 |
| R2506_S20_L001 | 48h with 100nM Rapa | HGPS - Fibroblasts | R1 | GGACTTGG-CGCAGACG | 3,103,306,911 | 30,725,811 | 96.63 | 36.47 |
| R2506_S20_L001 | 48h with 100nM Rapa | HGPS - Fibroblasts | R2 | GGACTTGG-CGCAGACG | 3,103,306,911 | 30,725,811 | 96.19 | 36.39 |
| R3074_S21_L001 | Control | HGPS - Fibroblasts | R1 | AAGTCCAA-TATGAGTA | 3,408,218,639 | 33,744,739 | 96.70 | 36.48 |
| R3074_S21_L001 | Control | HGPS - Fibroblasts | R2 | AAGTCCAA-TATGAGTA | 3,408,218,639 | 33,744,739 | 95.73 | 36.31 |
| R3075_S22_L001 | 48h with 12.5uM Pep14 | HGPS - Fibroblasts | R1 | ATCCACTG-AGGTGCGT | 3,467,352,523 | 34,330,223 | 96.58 | 36.46 |
| R3075_S22_L001 | 48h with 12.5uM Pep14 | HGPS - Fibroblasts | R2 | ATCCACTG-AGGTGCGT | 3,467,352,523 | 34,330,223 | 95.42 | 36.25 |
| R3076_S23_L001 | 48h with 100nM Rapa | HGPS - Fibroblasts | R1 | GCTTGTCA-GAACATAC | 3,238,628,731 | 32,065,631 | 96.65 | 36.47 |
| R3076_S23_L001 | 48h with 100nM Rapa | HGPS - Fibroblasts | R2 | GCTTGTCA-GAACATAC | 3,238,628,731 | 32,065,631 | 95.39 | 36.24 |
| R3310_S24_L001 | Control | HGPS - Fibroblasts | R1 | CAAGCTAG-ACATAGCG | 3,569,528,163 | 35,341,863 | 96.71 | 36.48 |
| R3310_S24_L001 | Control | HGPS - Fibroblasts | R2 | CAAGCTAG-ACATAGCG | 3,569,528,163 | 35,341,863 | 95.68 | 36.29 |
| R3311_S25_L001 | 48h with 12.5uM Pep14 | HGPS - Fibroblasts | R1 | TGGATCGA-GTGCGATA | 3,847,078,486 | 38,089,886 | 96.56 | 36.46 |
| R3311_S25_L001 | 48h with 12.5uM Pep14 | HGPS - Fibroblasts | R2 | TGGATCGA-GTGCGATA | 3,847,078,486 | 38,089,886 | 95.84 | 36.32 |
| R3312_S26_L001 | 48h with 100nM Rapa | HGPS - Fibroblasts | R1 | AGTTCAGG-CCAACAGA | 4,034,131,193 | 39,941,893 | 96.65 | 36.47 |
| R3312_S26_L001 | 48h with 100nM Rapa | HGPS - Fibroblasts | R2 | AGTTCAGG-CCAACAGA | 4,034,131,193 | 39,941,893 | 95.66 | 36.29 |
| R2897_R2903_S237_L004 | Control | Heathy - Fibroblasts 41 y | R1 | CGGACAAC-TCCGGATT | 2,838,845,986 | 28,107,386 | 96.45 | 36.44 |
| R2897_R2903_S237_L004 | Control | Heathy - Fibroblasts 41 y | R2 | CGGACAAC-TCCGGATT | 2,838,845,986 | 28,107,386 | 93.46 | 35.81 |
| R2898_S238_L004 | 24h with 12.5uM Pep14 | Heathy - Fibroblasts 41 y | R1 | ATATGGAT-CTGTATTA | 3,051,956,289 | 30,217,389 | 96.46 | 36.45 |
| R2898_S238_L004 | 24h with 12.5uM Pep14 | Heathy - Fibroblasts 41 y | R2 | ATATGGAT-CTGTATTA | 3,051,956,289 | 30,217,389 | 93.28 | 35.80 |
| R2899_S239_L004 | 24h with 100nM Rapa | Heathy - Fibroblasts 41 y | R1 | GCGCAAGC-TCACGCCG | 2,920,320,767 | 28,914,067 | 96.17 | 36.40 |
| R2899_S239_L004 | 24h with 100nM Rapa | Heathy - Fibroblasts 41 y | R2 | GCGCAAGC-TCACGCCG | 2,920,320,767 | 28,914,067 | 92.18 | 35.56 |
| R3031_S240_L004 | Control | Heathy - Fibroblasts 41 y | R1 | AAGATACT-ACTTACAT | 3,440,721,247 | 34,066,547 | 96.61 | 36.47 |
| R3031_S240_L004 | Control | Heathy - Fibroblasts 41 y | R2 | AAGATACT-ACTTACAT | 3,440,721,247 | 34,066,547 | 94.34 | 36.01 |
| R3032_S241_L004 | 24h with 12.5uM Pep14 | Heathy - Fibroblasts 41 y | R1 | GGAGCGTC-GTCCGTGC | 3,516,011,899 | 34,811,999 | 96.34 | 36.43 |
| R3032_S241_L004 | 24h with 12.5uM Pep14 | Heathy - Fibroblasts 41 y | R2 | GGAGCGTC-GTCCGTGC | 3,516,011,899 | 34,811,999 | 94.29 | 36.00 |
| R3033_S242_L004 | 24h with 100nM Rapa | Heathy - Fibroblasts 41 y | R1 | ATGGCATG-AAGGTACC | 4,036,709,521 | 39,967,421 | 96.64 | 36.48 |
| R3033_S242_L004 | 24h with 100nM Rapa | Heathy - Fibroblasts 41 y | R2 | ATGGCATG-AAGGTACC | 4,036,709,521 | 39,967,421 | 93.41 | 35.88 |
| R3301_S243_L004 | Control | Heathy - Fibroblasts 41 y | R1 | GCAATGCA-GGAACGTT | 3,051,587,235 | 30,213,735 | 96.38 | 36.43 |
| R3301_S243_L004 | Control | Heathy - Fibroblasts 41 y | R2 | GCAATGCA-GGAACGTT | 3,051,587,235 | 30,213,735 | 93.37 | 35.81 |
| R3302_S244_L004 | 24h with 12.5uM Pep14 | Heathy - Fibroblasts 41 y | R1 | GTTCCAAT-AATTCTGC | 4,096,673,120 | 40,561,120 | 96.53 | 36.46 |
| R3302_S244_L004 | 24h with 12.5uM Pep14 | Heathy - Fibroblasts 41 y | R2 | GTTCCAAT-AATTCTGC | 4,096,673,120 | 40,561,120 | 94.18 | 36.00 |
| R3303_S245_L004 | 24h with 100nM Rapa | Heathy - Fibroblasts 41 y | R1 | ACCTTGGC-GGCCTCAT | 2,925,069,888 | 28,961,088 | 96.48 | 36.45 |
| R3303_S245_L004 | 24h with 100nM Rapa | Heathy - Fibroblasts 41 y | R2 | ACCTTGGC-GGCCTCAT | 2,925,069,888 | 28,961,088 | 93.02 | 35.79 |
