## Supplementary material for "Senotherapeutic peptide reduces skin biological age and improves skin health markers": Sup Table 3

Supplementary Table 3 - Pathway Enrichment identified using the program Enrich with the top 20 genes modulated by Peptide14 as query genes. The KEGG 2019 database was used.

| **pathway** | **genes** | **pValue** | **adjPValue** |
| --- | --- | --- | --- |
| **Longevity regulating pathway** | RB1CC1 \| SIRT1 | 0.004609052918383906 | 1 |
| **FoxO signaling pathway** | RBL2 \| SIRT1 | 0.0075988978232358175 | 1 |
| **Signaling pathways regulating pluripotency of stem cells** | ID4 \| IL6ST | 0.008394292914981142 | 0.8618140726047306 |
| **Cellular senescence** | RBL2 \| SIRT1 | 0.010994232593906693 | 0.8465559097308153 |
| **Huntington disease** | NDUFS8 \| AP2S1 | 0.015702300024415403 | 0.9672616815039887 |
| **Viral carcinogenesis** | RBL2 \| IL6ST | 0.016953582060231374 | 0.8702838790918772 |
| **Mannose type O-glycan biosynthesis** | POMK | 0.02276098349039011 | 1 |
| **Endocytosis** | WASHC4 \| AP2S1 | 0.024373661102140152 | 0.9383859524323959 |
| **Fat digestion and absorption** | ABCA1 | 0.040229787776673645 | 1 |
| **ABC transporters** | ABCA1 | 0.044071254437806 | 1 |
| **Endocrine and other factor-regulated calcium reabsorption** | AP2S1 | 0.046942764384003306 | 1 |
| **Cholesterol metabolism** | ABCA1 | 0.04885254998202776 | 1 |
