## Supplementary material for "Senotherapeutic peptide reduces skin biological age and improves skin health markers": Sup Table 4

Supplementary Table 4 - Top 10 Pathways Enriched identified using the program Enrich with the top 90 genes modulated by Peptide14 as query genes. The KEGG 2019 database was used.

| **pathway** | **genes** | **pValue** | **adjPValue** |
| --- | --- | --- | --- |
| **Endocytosis** | EEA1 \| PSD3 \| AP2S1 \| RAB11FIP2 \| TGFBR1 | 0.0033006574257491645 | 1 |
| **TGF-beta signaling pathway** | PPP2R1A \| ID4 \| TGFBR1 | 0.00603936607241716 | 0.9300623751522427 |
| **Th17 cell differentiation** | IL6ST \| HIF1A \| TGFBR1 | 0.009709442825416097 | 0.9968361300760527 |
| **FoxO signaling pathway** | RBL2 \| SIRT1 \| TGFBR1 | 0.0170418132441655 | 1 |
| **D-Glutamine and D-glutamate metabolism** | GLUD2 | 0.020334452124159075 | 1 |
| **Parkinson disease** | COX8A \| NDUFS8 \| PARK7 | 0.020642762158158173 | 1 |
| **Cellular senescence** | RBL2 \| SIRT1 \| TGFBR1 | 0.02809925527763663 | 1 |
| **Renal cell carcinoma** | VHL \| HIF1A | 0.03263286994541969 | 1 |
| **Adherens junction** | YES1 \| TGFBR1 | 0.0352761374472165 | 1 |
| **Huntington disease** | COX8A \| NDUFS8 \| AP2S1 | 0.04499446485011267 | 1 |
