## Supplementary figures and images for "Senotherapeutic peptide reduces skin biological age and improves skin health markers"

### Sup Fig 1

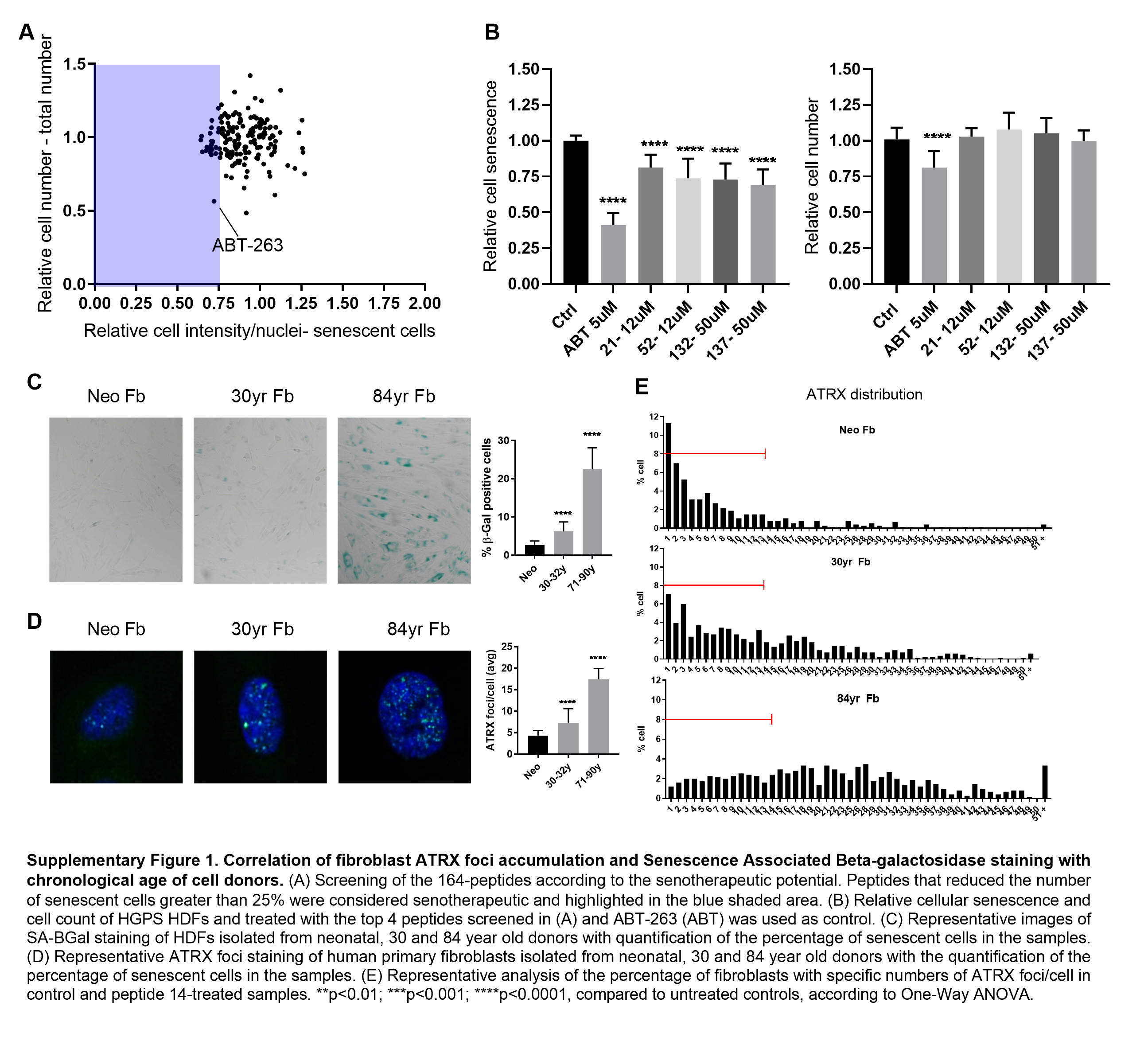

### Sup Fig 2

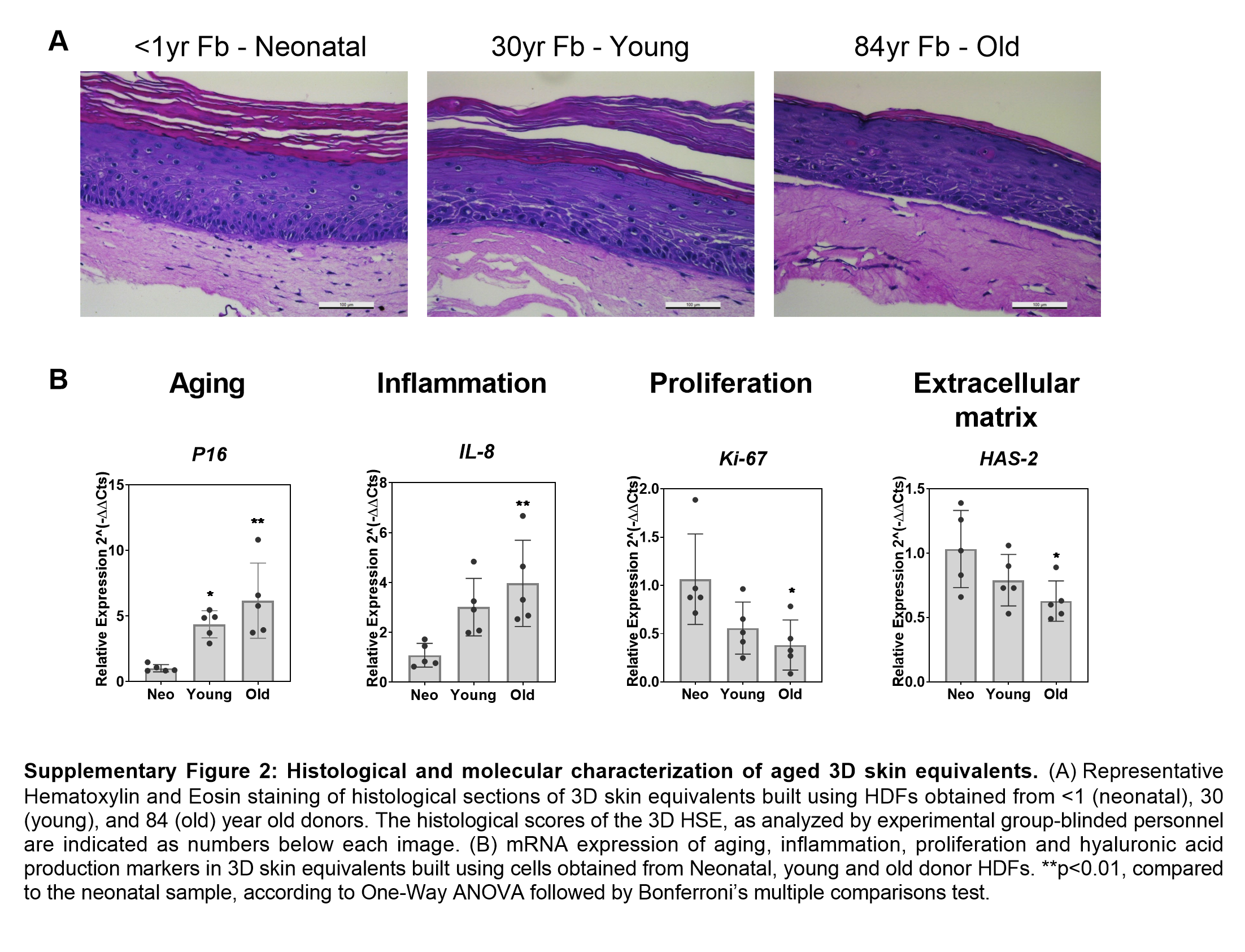

### Sup FIg 3

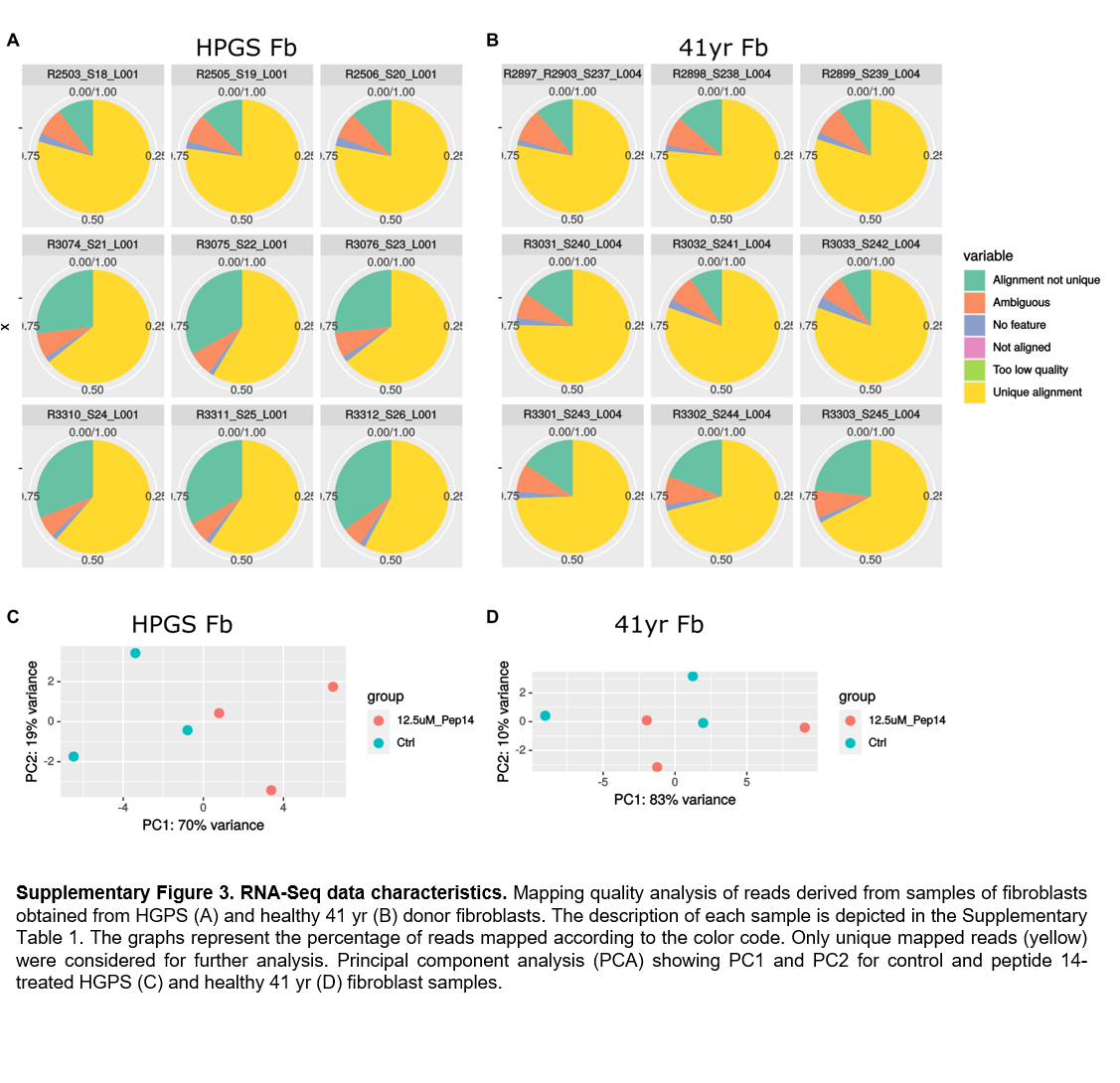

### Sup Fig 4

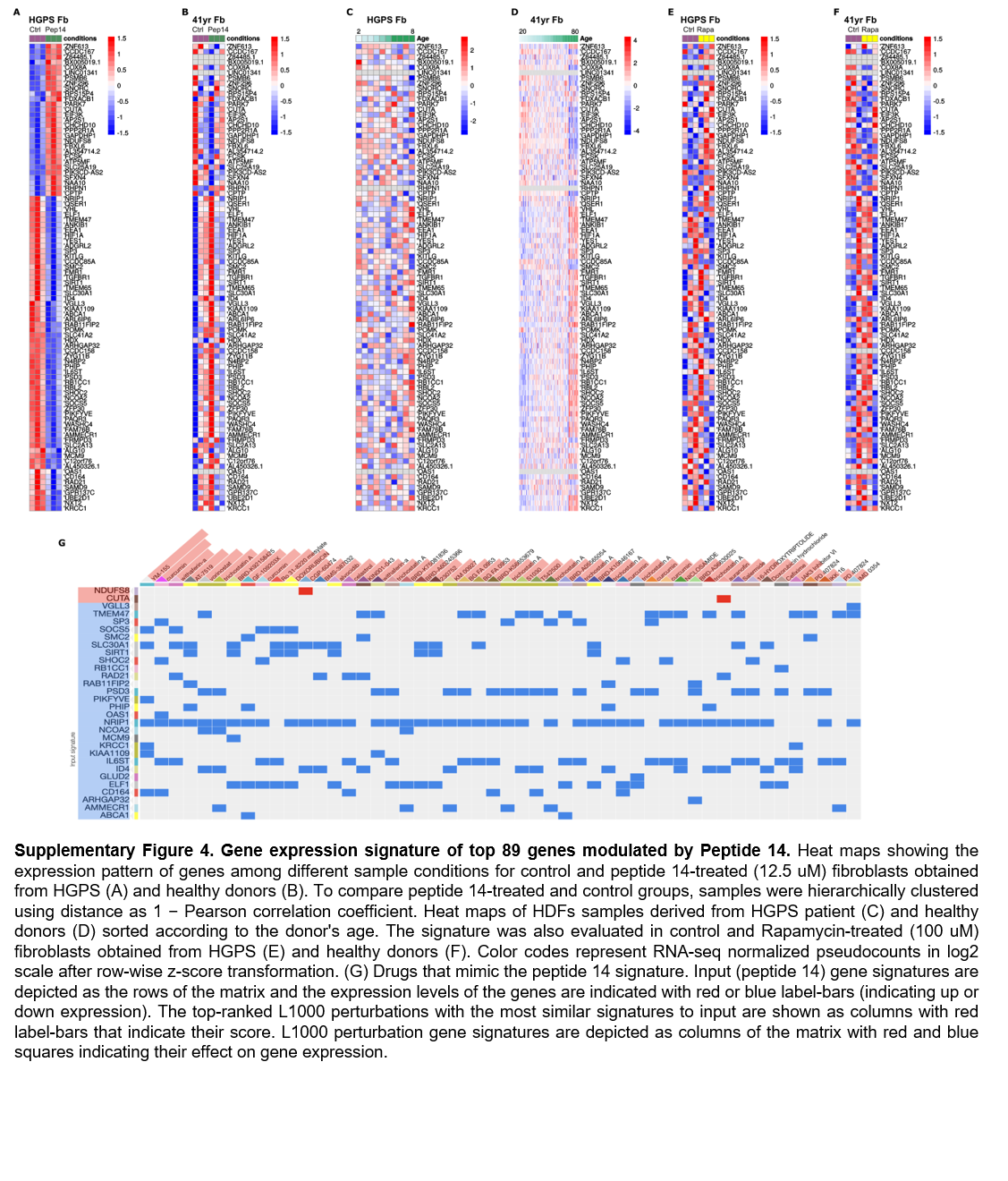
